# Tumour-derived LAMA5 is critical for tumour initiation and controls progression and phenotype in luminal breast cancer

**DOI:** 10.1101/2024.12.26.630367

**Authors:** Alexandra Ritchie, Priyal Dhawan, Eveliina Kailari, Jere Linden, Leander Blaas, Valerio Izzi, Pekka Katajisto, Johanna I. Englund

## Abstract

Basement membrane (BM) supports and regulates the structural integrity, function and differentiation of epithelial tissues and protects against breast cancer invasion to stroma. Here we show that the BM component LAMA5 is critical for initiation of luminal mammary tumours, and controls tumour progression and phenotype development. LAMA5 is overexpressed in human breast carcinomas and *LAMA5* downregulation attenuates growth of human breast cancer cells. Prepubertal luminal deletion of *Lama5* in MMTV-PyMT mice results in marked reduction in emergence of early hyperplasias, along with a shift towards luminal progenitor-like phenotype. However, single allele deletion, but not biallelic deletion of *Lama5* inhibits growth and progression towards advanced mammary carcinomas. *Lama5* deficient luminal epithelial cells collectively display widespread alterations in Fibroblast growth factor (Fgf) signalling genes, including increased expression of Fgf receptor 2 (Fgfr2). Inhibition of Fgf receptors decreases growth and induces apoptosis also in biallelic-*Lama5-*deleted organoids without affecting wildtype organoids. Our results demonstrate a critical role for the BM component LAMA5 in mammary tumour initiation and reveal mechanisms of ECM-epithelial interplay in breast tumour progression and phenotype maintenance.

## INTRODUCTION

Cancer is a disease characterized by uncontrolled cell growth and invasion into surrounding tissue. The basement membrane (BM) is a specialized, sheet-like form of extracellular matrix (ECM) separating epithelial cells from the stroma ^1^. It is widely considered a protective element against tumour progression from ductal carcinoma in situ (DCIS) to invasive breast cancer ^1–3^, and discontinuity of the BM sheet and degradation or loss of BM components are frequently reported in invasive tumours ^4^.

In healthy tissue, epithelial adhesion to the BM plays a critical role in maintaining structural integrity and regulating cell behaviour, including proliferation, differentiation, and migration ^1^. BM sheets are typically composed of collagen IV and laminin networks and attached glycoproteins ^4,5^. Laminins are heterotrimeric proteins comprising of α, β and γ subunits essential for BM formation and expressed in tissue-specific and temporally regulated patterns ^4^. In healthy mammary epithelium the major laminin trimers expressed are laminin-111 (composed of α1, β1 and γ1 subunits), laminin −332 and laminin −511/521 trimers ^6,7^. We have previously demonstrated that laminin α5 (found in laminin-511/521) is produced predominantly by the hormone receptor positive (HR+) luminal epithelial cells and adhesion to laminin α5-containing laminin trimers is critical for the mammary epithelial growth during puberty and pregnancy via supporting HR+ cell differentiation, luminal gene expression, and luminal-basal signalling^8,9^.

Despite the tumour-protective barrier function of BM, certain BM components, including laminin subunits α4 and α5, laminin-332, agrin, and nidogen are found overexpressed in human breast tumours as well as in other malignancies such as prostate, lung, and liver carcinomas ^10–16^. A recent study furthermore showed that mammary tumour cells actively produce and assemble BM during tumour development ^17^. Moreover, BM constituents, such as laminin-332 and laminin-511 increase cell migration *in vitro* and are utilized by tumour cells for motility and metastatic spread *in vivo* ^10,15,18^ suggesting that they contribute to tumour progression separately from the protective tissue barrier function of BM. While stromal cells produce most of the tumour ECM, recent studies have elucidated the importance of tumour-cell derived ECM in tumour progression and as indicator of patient survival ^12^.

Considering the contribution of BM components to tumour progression, and the essential role of luminally derived laminin α5 in the formation of healthy mammary epithelium, we sought to explore whether laminin α5 contributes to initiation and progression of luminal tumours. Here we show that laminin α5 is abundantly expressed especially in luminal mammary tumours and that its loss inhibits growth of human luminal breast cancer cells *in vitro* and markedly reduces early tumour initiation and alters tumour progression in mouse. Furthermore, loss of Lama5 results in an epithelial phenotype switch towards an immature luminal progenitor-like phenotype along with widespread changes in Fibroblast growth factor (Fgf) signalling in a luminal tumorigenesis mouse model *in vivo.* Our results reveal a critical novel role for a BM component laminin α5 in mammary tumour initiation and demonstrate the profound effects of altered tumour ECM composition for tumour phenotype and breast tumour progression.

## RESULTS

### *LAMA5* is overexpressed in human luminal breast carcinomas

We first sought to determine *LAMA5* expression pattern in normal human breast tissue and in human breast tumours by performing RNA in situ hybridization (ISH) for *LAMA5* on a tissue micro array (TMA) set with 100 invasive human breast carcinoma cases and 9 samples of adjacent normal breast tissue.

Normal breast tissues exhibited a luminal staining pattern of *LAMA5* ISH (**Fig 1a, Fig S1a**) similarly to mouse mammary tissue^8^. The staining was observed in majority of the luminal cells in both ducts and terminal ductal lobular units (TDLU) (**Fig S1b**). The breast cancer tissues on the other hand showed variable staining patterns ranging from weak to very strong (**Fig 1b**). To quantitate the differences between the normal and tumour samples, we calculated the intensity and coverage of the ISH staining from the TMA spots using our image analysis software Tonga^19^. Tumour spots exhibited on average 5.9 times more *LAMA5* ISH staining than normal breast tissue (mean normal 2.13 ‰ ± 2.3 σ, tumour 12.58 ‰ ± 16.34 σ) (**Fig 1b**).

**Figure 1.**
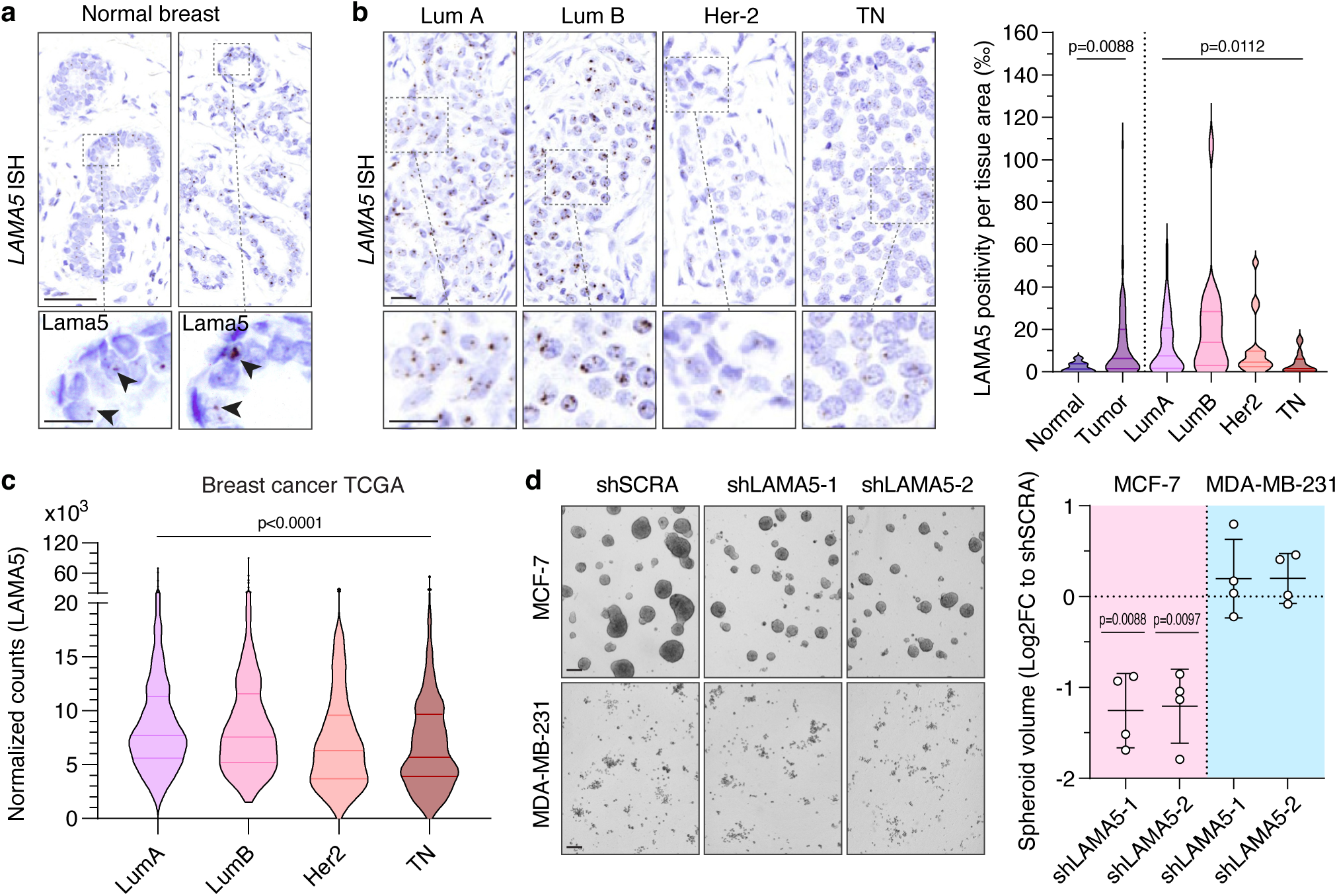
LAMA5 is overexpressed in human luminal breast tumours and affects growth of luminal breast cancer cells. **a)** In situ hybridization (ISH) of *LAMA5* and negative control probe in normal human breast. Scale bar 50 μm or 10 µm (zoom ins). **b)** Representative images of ISH of *LAMA5* in different breast tumour subtypes including luminal A (LumA), luminal B (Lum B), Her-2 positive (Her-2) and triple negative (TN). Scale bar 20 µm. Right graph shows quantiation of *LAMA5* positivity parts per thousand (‰) per tissue area in TMA samples of normal breast tissue and breast tumors (left). Right side demonstrates *LAMA5* positivity when same samples are divided into molecular subtypes. Normal n=19, LumA n=46, LumB n=20, Her-2 n=17, TN n=17. Statistical analysis is performed using a Kruskal-Wallis test (tumour subtypes) and a Mann-Whitney test (tumour vs normal). **c)** Normalized LAMA5 gene expression in differrent breast cancer subtypes from The Cancer Genome Atlas. LumA n=444 LumB n=283, Her-2 n=72, TN n=116. Statistical analysis is performed using one-way ANOVA (Kruskal-Wallis test). d) Representative images of MCF-7 and and MDA-MB 231 cells expressing SCRA or LAMA5 shRNA grown as spheroids in low-adhesion culture for 10 days. Scale bar 200 μm. Right graph shows quantitation of spheroid growth as log fold change (logFC) compared to control (shSCRA). Each point rerpesent individual experiment. Statistical analysis is performed using one-sample t tests against zero.

We also investigated whether *LAMA5* expression correlates with specific breast cancer subtypes by categorizing the tumour samples by molecular subtype, estrogen receptor (ER) status, and grade. Luminal B (Lum B, n=20 samples) subtype exhibited on average the strongest *LAMA5* ISH staining, while in Luminal A (Lum A, n=46 samples) and Her-2 positive (Her-2, n=17 samples) showed intermediate and triple-negative subtype (TN, n=17 samples) the lowest level of staining (**Fig 1b**). No correlation to ER positivity or grade was observed (**Fig S1c**) most likely due to the small sample size of the TMA.

We additionally queried *LAMA5* expression from The Cancer Genome Atlas (TCGA) Breast Invasive Carcinoma cohort (TCGA-BRCA) by molecular subtype (n=915) and observed highest *LAMA5* expression in the Lum B and Lum A subgroups and lowest in the TN subgroup (**Fig 1c and S1d**), similarly to our ISH data. Moreover, higher *LAMA5* expression was associated with advanced tumour stages (n=887) II and III in Lum A tumours. (**Fig S1d**). Taken together, our ISH data and published expression data suggests that there is a robust overexpression of *LAMA5* in human breast cancers especially in luminal subtypes.

### *LAMA5* downregulation decreases growth in luminal breast cancer cells

As overexpression of *LAMA5* was most noticeable in luminal subtypes as opposed to the triple-negative subtype, we hypothesized that LAMA5 could give growth advantage specifically to luminal breast cancer cells. To test this, we downregulated *LAMA5* from luminal MCF-7 and triple-negative MDA-MB-231 cell lines using two independent shRNA hairpins (shLAMA5-1 and −2) and achieved ∼50% reduction in LAMA5 protein deposition with both constructs compared to control scramble shRNA (shSCRA) (**Fig S1e-g)**. Using the cell lines, we first investigated the effect of *LAMA5* downregulation to 2D cell growth using live cell imaging. MCF-7 cells expressing shLAMA5-1 and shLAMA5-2 grew approximately 25% and 30% less than the shSCRA cells, with doubling time increased from 28 hours of shSCRA to 34 hours and 35 hours respectively (**Fig S1h**). In MDA-MB-231 cells however, LAMA5 silencing did not alter growth and doubling time (**Fig S1h**). EdU incorporation was reduced by 19% in LAMA5 silenced MCF-7 cells while MDA-MB-231 cells were unaffected, suggesting that *LAMA5* promotes MCF7 cell growth via proliferation (**Fig S1i**).

Since 2D monolayer culture only partially models’ growth capacity of breast cancer cells, we assayed the growth of *LAMA5* downregulated cells in 3D suspension culture environment ^20^. MCF-7 cells grew in roughly symmetrical spheroids, while MDA-MB-231 cells rather formed loosely connected clusters or sheet-resembling structures (**Fig 1d**). After 10 days in culture, the total growth of MCF-7 shLAMA5-1 and shLAMA5-2 cells was less than half compared to shSCRA, while no differences in total growth were seen in MDA-MB-231 cells (**Fig 1d**). We additionally immunostained LAMA5 in the final day 10 spheroids or clusters confirming a comparable decrease of LAMA5 protein as in our original analyses (**Fig S1f-g**). Taken together, our data demonstrates that downregulation of *LAMA5* results in proliferative growth decline in luminal but not in triple-negative breast cancer cells both in 2D monolayer and 3D suspension cultures.

### Tumour-derived LAMA5 is critical for mammary tumour initiation

As our human data showed LAMA5 overexpression to confer functional growth advantage for luminal cancer cells *in vitro*, we next investigated the role of Lama5 in tumorigenesis *in vivo*. We utilized the polyoma middle-T (PyMT) transgenic mouse model ^21^, where female MMTV-PyMT animals develop spontaneous luminal lesions starting from hyperplasia at 4-6 weeks of age that progress through adenoma/mammary intraepithelial neoplasia to carcinoma around 8-14 weeks of age recapitulating the progression of human luminal tumours ^21,22^.

We first assayed whether expression of *Lama5* or other major BM laminin α chains produced by mammary epithelium is altered also in MMTV-PyMT tumorigenesis. We thus compared expression of *Lama1*, *Lama3* and *Lama5* in normal mammary epithelium, early *in situ* lesions representing hyperplasia or early adenoma (hereafter simplified as hyperplasia), and invasive adenocarcinomas, by RNA ISH (**Fig 2a-b, Fig S2a**). Staining for *Lama1* and *Lama3*, which are mostly expressed by basal MECs in normal epithelium (**Fig 2a**), was observed in low levels in both hyperplasia lesions and adenocarcinomas (**Fig 2a-b**). While *Lama3* expression was present mostly intermittently on the edges of tumour lobules, likely representing remaining basal-like cells, *Lama1* was expressed by the tumour mass (**Fig 2a**). Although mean epithelial staining for both *Lama1* and *Lama3* decreased in both hyperplasia and adenocarcinoma in comparison to normal tissue, the change was statistically significant only for *Lama3*, likely due to small sample size (n=4). Also, *Lama5* staining showed a major decrease in hyperplasia lesions in comparison to normal ducts (**Fig 2a-b**), likely due to hyperplasia containing less differentiated luminal cells, which express *Lama5* in high amounts in normal epithelium (Fig 2a and ^8^). Interstingly however, *Lama5* staining was found within the lesions, and a clear increase in *Lama5* staining was observed in progression from hyperplasia to adenocarcinoma, (**Fig 2a-b**). Together with our data from human tumours (**Fig 1**), this suggests tumour-derived *Lama5* and high *Lama5* expression to be common features of advanced luminal mammary tumours, contrary to other laminin α chains.

**Figure 2.**
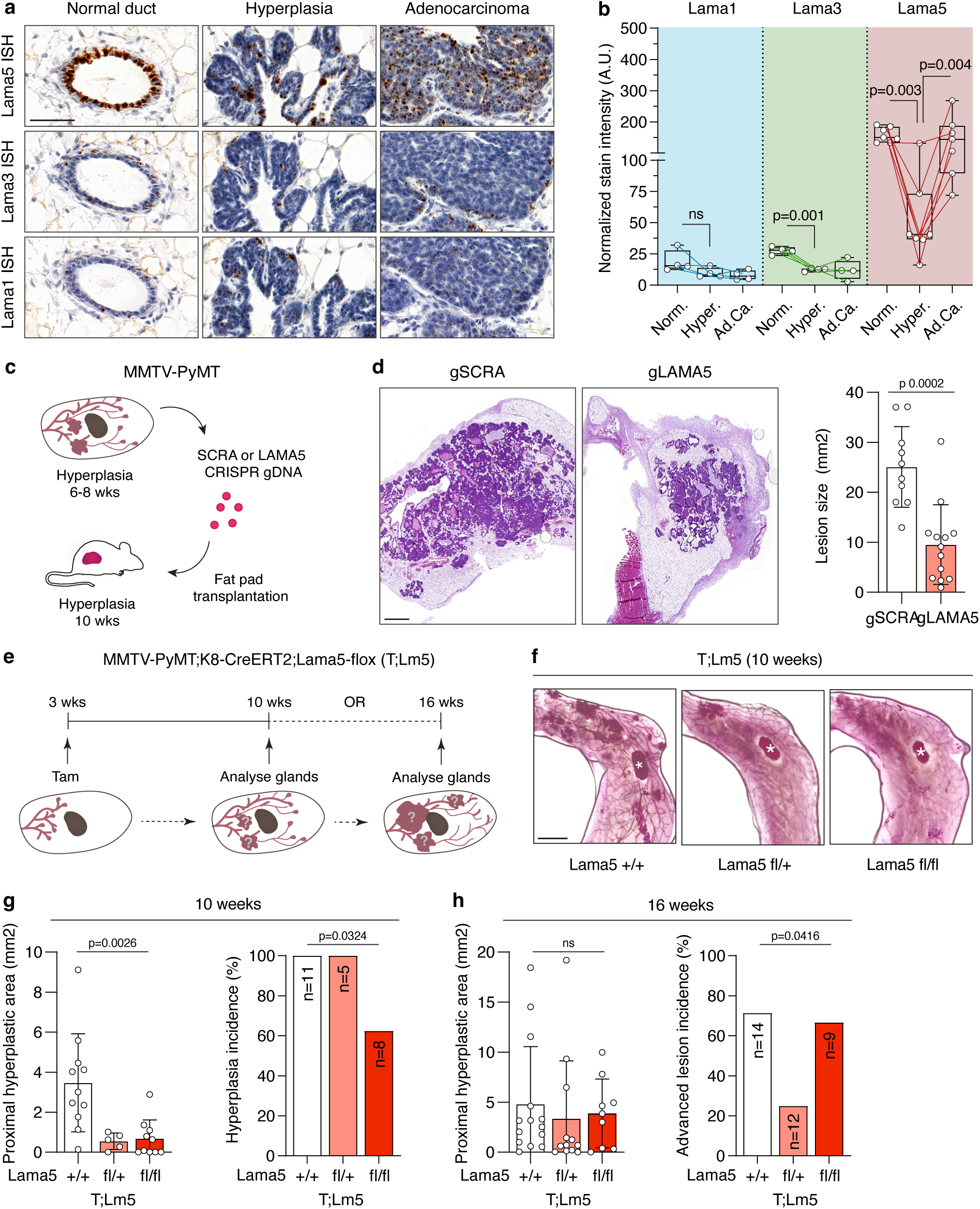
Tumour-derived Lama5 is critical for early tumour growth. **a)** Representative images of normal ducts, hyperplasias and adenocarcinomas from MMTV-PyMT mice stained with Lama1, Lama3, or Lama5 ISH. Scale bar 50 µm. **b)** Quantiation of Lama1, Lama3 and Lama5 positivity in normal ducts (Norm.) hyperplasias (Hyper.) and adenocarcinomas (Ad.Ca.) in MMTV-PyMT mice. Each spot represents 1 mouse analysed. Lines represents matching of samples from the same mouse. Matched t-tests for ratios were used for statistical testing. **c)** Schematic of the experimental set-up, where mammary epithelial cells from 6-8 week old MMTV-PyMT are isolated and either LAMA5 or SCRA gDNA is introduced to cells, and the cells are transplanted into wild type hosts. Mammary glands are collected 6 weeks later when the recipient mice were 10 weeks old and resulting growth is analysed. **d)** Representative HE images of mammary glands transplanted with gLAMA5 or gSCRA expressing cells as described in c. Scale bar 1 mm. Graph shows quantitation of lesion size, each dot represents one gland analysed. Statistical analysis was performed using two-tailed Student’s t-test. **e)** Schematic of the experimental set-up of MMTV-PyMT;K8-CreERT2;Lama5-flox (T;Lm5) and control mice. **f)** Representative carmine-alumn stained mammary glands of 10-week-old T;Lm5 mice, asterisk denotes the lymph node. Scale bar 2 mm. **g)** Lesion size and lesion incidence in the 10-week-old T;Lm5 mice. Each dot represents one gland analysed. Kruskal-Wallis test and a Chi-squared test were used for statistical testing, respectively. **h)** Lesion area and advanced lesion incidence in the 16-week-old T;Lm5 mice.

To examine whether *Lama5* loss attenuates tumour cell growth in MMTV-PyMT tumours, we isolated mammary epithelial cells from MMTV-PyMT mice at 6-8 weeks of age when hyperplastic nodules have started appearing, but prior to increase of *Lama5* expression based on our ISH data, and deleted *Lama5* from the cells with CRISPR silencing (**Fig S2b** and ^8^). Using 3D organoid culture, we observed a clear decrease in size of *Lama5*-targeted organoids (*Lama5* gRNA, gLAMA5) after 7 days in culture in comparison to scrambled gRNA (gSCRA) controls, suggesting a growth reduction also in mouse mammary tumour cells (**Fig S2c**).

To assay growth in an *in vivo* environment we next transplanted the *Lama5*-targeted and control cells into cleared mammary fat pads of wildtype recipient mice and analysed the glands 6 weeks later (**Fig 2c**). The outgrowths observed in the glands were mostly hyperplastic lesions, with some normal-like ductal growth observed specifically in the *Lama5*-silenced samples (**Fig 2d, FigS2d**). When measured, we observed a significant reduction in the lesion size in *Lama5*-targeted outgrowths compared to controls (**Fig 2d**). Immunohistochemical (IHC) analysis of the outgrowths showed no difference in neither the proliferation marker Ki-67 (**Fig S2e**), nor apoptosis as measured by TUNEL positivity (**Fig S2e**). These results suggest that the reduction in hyperplastic lesion size in transplants with silenced *Lama5* resulted from reduced initial tumour growth, rather than loss of proliferative growth or increased cell death.

To test this further, we crossed the MMTV-PyMT mice with Lama5 flox ^23^ and K8-CreERT2 ^24^ mice (hereafter T;Lm5) to allow targeted *Lama5* deletion specifically in the tumour initiating luminal MECs. Using these animals, we determined whether early loss of *Lama5* affects initiation of hyperplastic lesions during pubertal growth between 6-8 weeks of age, by injecting tamoxifen into prepubertal 3-week-old T;Lm5fl/fl, T;Lm5fl/+ and T;Lm5+/+ control mice and analysing the mammary glands at 10 weeks of age (**Fig 2e**). Wholemounts of the control mammary glands exhibited frequent hyperplastic nodules, whereas *Lama5*-deleted glands exhibited mainly normal ducts with fewer hyperplastic lesions (**Fig 2f**). In accordance with our earlier findings^8^, also the ductal network in T;Lm5fl/fl epithelium was underdeveloped, yet the proximal ductal growth and density remained unaltered (**Fig S2f-g**). However, when measuring the total proximal hyperplastic lesion area, we observed a notable 5-fold decrease in both T;Lm5fl/+ and T;Lm5fl/fl glands in comparison to control glands (**Fig 2g**). Furthermore, while all T;Lm5fl/+ and control mice developed at least one hyperplastic lesion, 40% of T;Lm5fl/fl glands were completely free of visible hyperplasia (**Fig 2g)**. We additionally quantitated Ki-67 positivity in the hyperplastic lesions to measure proliferation but did not observe significant differences between genotypes (**Fig S2h**), suggesting that early deletion of *Lama5* from the luminal epithelial cells restrains PyMT-oncogene-induced tumour initiation without affecting tumour cell proliferation, similarly to our transplant experiment.

To determine whether prepubertal *Lama5* loss can also restrain further tumour development and prevent progression from hyperplastic nodules towards large adenomas and early carcinomas typically developing between 8 and 14 weeks of age ^21,22^, we analysed mammary glands of tamoxifen-injected 3-week-old T;Lm5fl/fl, T;Lm5fl/+ and T;Lm5+/+ control mice at 16 weeks of age (**Fig 2e**). The control mammary glands exhibited numerous hyperplastic nodules similarly to the 10-week-old mice, along with single advanced lesions above 2 mm^2^ in size (**Fig S2i**). However, no difference in the total proximal lesion area was seen between the *Lama5*-deficient glands and the controls (**Fig 2h**). We also quantitated the incidence of the advanced single lesions (>2 mm^2^), which were interestingly less frequent in T;Lm5fl/+ mice compared to control and T;Lm5fl/fl mice, which both showed similar advanced lesion incidence (**Fig 2h**). Altogether our data suggests that deletion of *Lama5* critically affects tumour initiation by delaying hyperplasia formation, while tumour progression towards advanced lesions was reduced only by deletion of one *Lama5* allele. This raises the possibility that early biallelic *Lama5* deletion eventually induces compensatory mechanisms leading to restoration of tumour growth and progression despite initial delay in tumour formation.

### Biallelic deletion of *Lama5* selects for LAMA5-independent growth during tumour progression

Next, we investigated whether *Lama5* deletion can modulate the progression of early hyperplastic lesions towards adenocarcinomas when deleted later in the already established hyperplasias. We injected tamoxifen into T;Lm5fl/fl, T;Lm5fl/+ mice and T;Lm5+/+ control mice at the age of 8 weeks when established hyperplasias are present and typically start progressing towards carcinomas, and analysed the mice when first tumour with 1 cm diameter was observed by palpation (**Fig 3a**). Mean latency to appearance of the first tumour in the control mice was 109±32 days (95±39 in mice not treated with tamoxifen), which was comparable to T;Lm5fl/+ (112±41 days) and T;Lm5fl/fl mice (100±27 days; **Fig 3b**). Also, the average tumour size was similar in all groups, as expected due to the common tumour growth end point (**Fig 3c**). However, loss of one *Lama5* allele led to a significant reduction in the tumour multiplicity with a mean of 1,75 palpable tumours per animal compared to 5,25 and 4,6 in T;Lm5+/+ and no tamoxifen control groups, respectively. Unexpectedly, no reduction was observed in biallelic-deleted T;Lm5fl/fl animals, with mean of 4,75 tumours per animal (**Fig 3c**). An additional Ki-67 positivity analysis revealed also decreased cell proliferation in T;Lm5fl/+ tumours, but not in T;Lm5fl/fl tumours (**Fig 3d**), suggesting slower growth as a result from partial *Lama5* loss, while biallelic-*Lama5-*deleted tumours remain comparable to controls. This supports the possibility of biallelic-*Lama5-*deleted tumours acquiring compensatory LAMA5-independent growth mechanisms during tumour progression, irrespectively whether *Lama5* loss occurs before or after initial hyperplasia formation.

**Figure 3.**
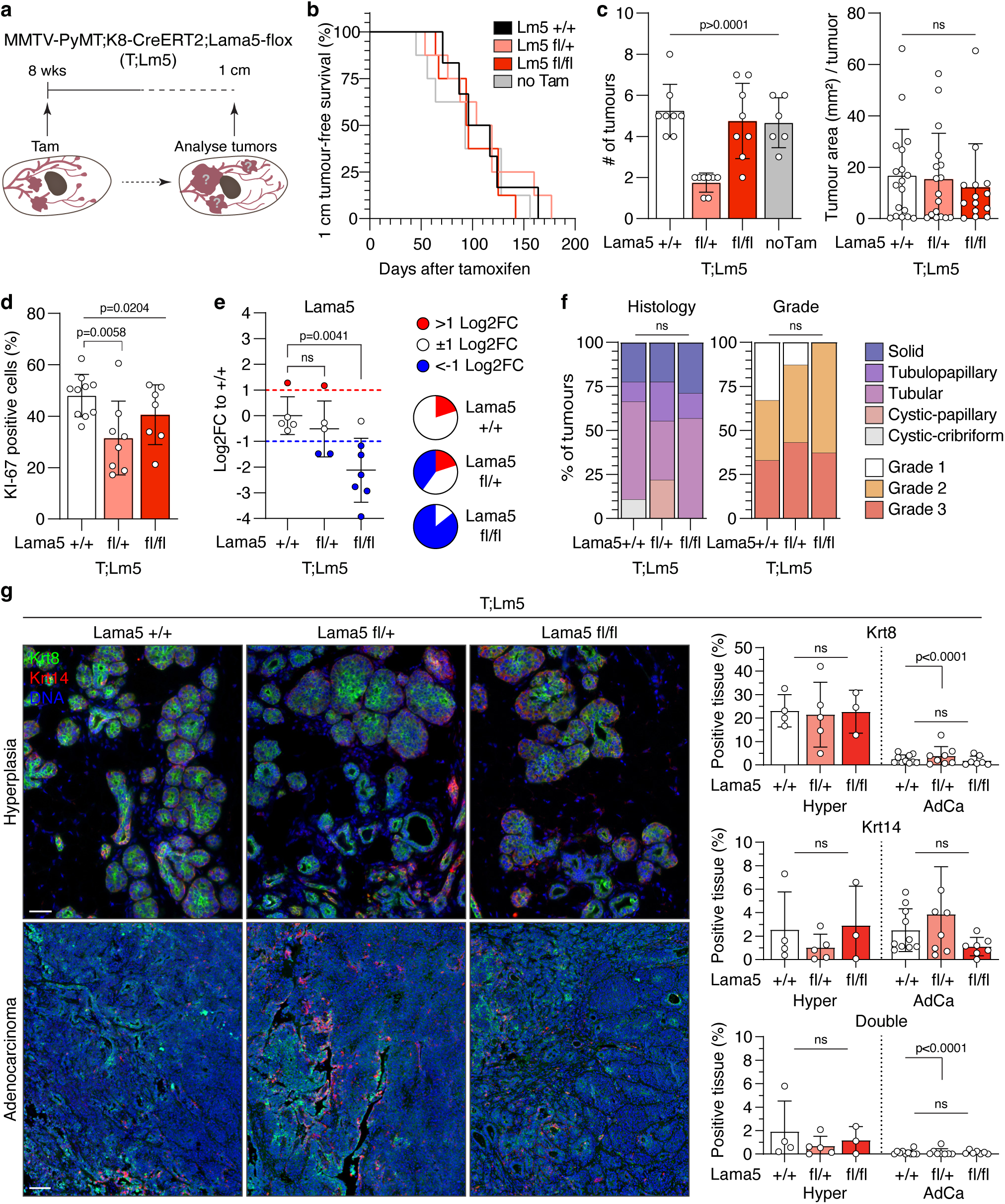
Biallelic Lama5 deletion in adenocarcinomas selects for LAMA5-independent growth without altering the tumour subtype. **a)** Schematic of the experimental set-up of MMTV-PyMT;K8-CreERT2;Lama5-flox (T;Lm5) and control mice. Tam = tamoxifen. **b)** Kaplan-Meier curve showing the probability of survival until appearance of the first 1 cm tumour in each genotype and MMTV-PyMT;Lama5 +/+ treated with vehicle control (no Tam). **c)** Quantiation of tumour area and multiplicity of tumours in each genotype and in no Tam control mice. Each spot represents 1 tumour analyzed. Statistical analysis is performed using two-way ANOVA with Tukey’s multiple comparison test. **d)** Analysis of Ki-67 positivity in the T;Lm5 and control tumours. Each spot represents 1 tumour analyzed. Statistical analysis is performed using two-way ANOVA and a Fisher’s LSD test. e) qPCR analysis of Lama5 wild type (WT) allelle in the adenocarcinomas relative to GAPDH. Each dot represents 1 carcino-ma analyzed. Red dots represent carcinomas with >1 Log2 fold change and blue dots with <-1 Log2 fold change of expression in comparison to the control. Parts of whole charts show the frequency of carcinomas with the indicated expression level. One sample t-tests against 0 were used for statistical analysis. **f)** Frequency of different histological types and tumour grades in the T;Lm5 and control tumours (+/+ n=9, fl/+ n=9, fl/fl n=8). Statistical testing was performed using Chi-squared tests. g) Representative IF images of T;Lm5 hyperplasia (Hyper) with prepubertal Lama5 deletion and adenocarcinoma (AdCa) tumours with late Lama5 deletion stained with Krt8 (green) and Krt14 (magenta) antibodies. Scale bar 50 μm (hyper) and 100 μm (AdCa). Graphs show quantiation of Krt8 and Krt14 positivity, as well as double positivity (Krt8 and Krt14) in the lesions. Statistical analysis is performed using two-way ANOVA. Each spot represents 1 tumour analysed.

To exclude the possibility of T;Lm5fl/fl tumours having escaped recombination and therefore emerging in higher numbers compared to T;Lm5fl/+ tumours, we next analysed the expression of functional wild type (WT) Lama5 allele by qPCR ^8^. All T;Lm5fl/fl tumours expressed WT *Lama5* mRNA in greatly decreased levels (<-1 Log2FC to control mean) except for one tumour analysed. One T;Lm5fl/+ tumour was also found to express WT mRNA at high levels (>1 Log2FC of control mean), while all the other analysed T;Lm5fl/+ tumours expressed WT mRNA in levels below control mean (**Fig 3e**). These data indicate that while individual tumours may escape the *Lama5* deletion, majority of the tumours arising in T;Lm5fl/fl animals have likely acquired alternative, LAMA5-independent growth mechanisms.

### *Lama5* loss does not alter tumour histopathology or induce a basal-like conversion of luminal tumours

Distinct progression kinetics in T;Lm5fl/fl tumours irrespective of deletion timepoint led us to hypothesize that biallelic *Lama5* loss could predispose to tumours with altered characteristics. To test this, we classified the tumours as solid, tubulopapillary, tubular, cystic-papillary or cystic-cribriform subtypes based on previously described histopathological criteria ^25^, and graded them 1-3 by cell abnormality by adapting a grading system based on Elston and Ellis numeric method for human breast cancer ^26,27^ (**Table S1**). As shown previously in MMTV-PyMT tumours ^28^, we observed variations in grade and subtype. However, except for the majority of the tumours being of tubular subtype in T;Lm5+/+ and T;Lm5fl/fl tumours, and the absence T;Lm5fl/fl tumours of grade 1, the distributions of subtypes and grades were similar in all groups (**Fig 3f, Fig S3a**), indicating no significant changes in the MMTV-PyMT tumour histopathology upon *Lama5* loss to explain the altered progression kinetics of T;Lm5fl/fl tumours.

Luminal mammary tumour cells have been shown in specific circumstances to acquire basal characteristics, suggested to underlie emergence of the more aggressive basal-like subtypes ^29–31^. As we have previously shown LAMA5 to be necessary for luminal characteristics and HR+ differentiation in normal MECs ^8,9^ we next hypothesized that biallelic *Lama5* loss could trigger a HR+ luminal phenotype loss and conversion towards a basal-like phenotype in the MMTV-PyMT tumours, which are prone to losing their HR+ luminal characteristics upon progression to carcinoma^21^, explaining their altered growth pattern. To address this, we analysed expression of luminal and basal lineage markers KRT8 and KRT14 in the T;Lm5 adenocarcinomas with late *Lama5* deletion (**Fig 3a**), in the 10-week-old T;Lm5 hyperplastic lesions with prepubertal *Lama5* deletion (**Fig 2e**), as well as in the fat-pad-transplanted tumours where the outgrowths originate fully from the transplanted cells with CRISPR-mediated *Lama5* deletion. However, except for expression of KRT8 being more prominent in hyperplastic lesions in comparison to adenocarcinomas, we discovered no differences (**Fig 3g, Fig S3b**). We additionally analysed possible emergence of KRT8/KRT14 co-expressing double positive cells, which have previously been suggested to precede luminal-basal lineage conversion^29,32^ but observed no differences (**Fig 3g, Fig S3b**). Finally, we performed qPCR of *Lama5* low T;Lm5 adenocarcinomas with late *Lama5* deletion (**Fig 3a**, **Fig 3e**) with luminal markers *Krt8* and *Krt18* and basal markers *Krt5*, *Krt14* and *Lgr5* to address gene-level expression (**Fig S3c**) but also discovered no differences between the genotypes in any of the markers. Altogether, neither change in histopathology or in luminal-basal phenotype conversion explain increased tumour growth in the biallelic-*Lama5*-deleted tumours but rather suggest other underlying changes following *Lama5* loss.

### *Lama5* loss induces a shift towards progenitor-like phenotype

To understand the changes underlying the different growth patterns of single-allele-deleted and biallelic-*Lama5*-deleted tumours, we next carried out single cell RNA sequencing (scRNA-seq) of mammary epithelium and associated cells from 10-week-old mice with all genotype combinations of *Lama5* and MMTV-PyMT (**Fig 4a**). FACS sorted, hematopoietic lineage (Lin) negative (CD45-,CD31-,Ter119-) cells (**Fig S4a**) from altogether 5 mice with MMTV-PyMT were sequenced using 10X Chromium platform^33^.

**Figure 4.**
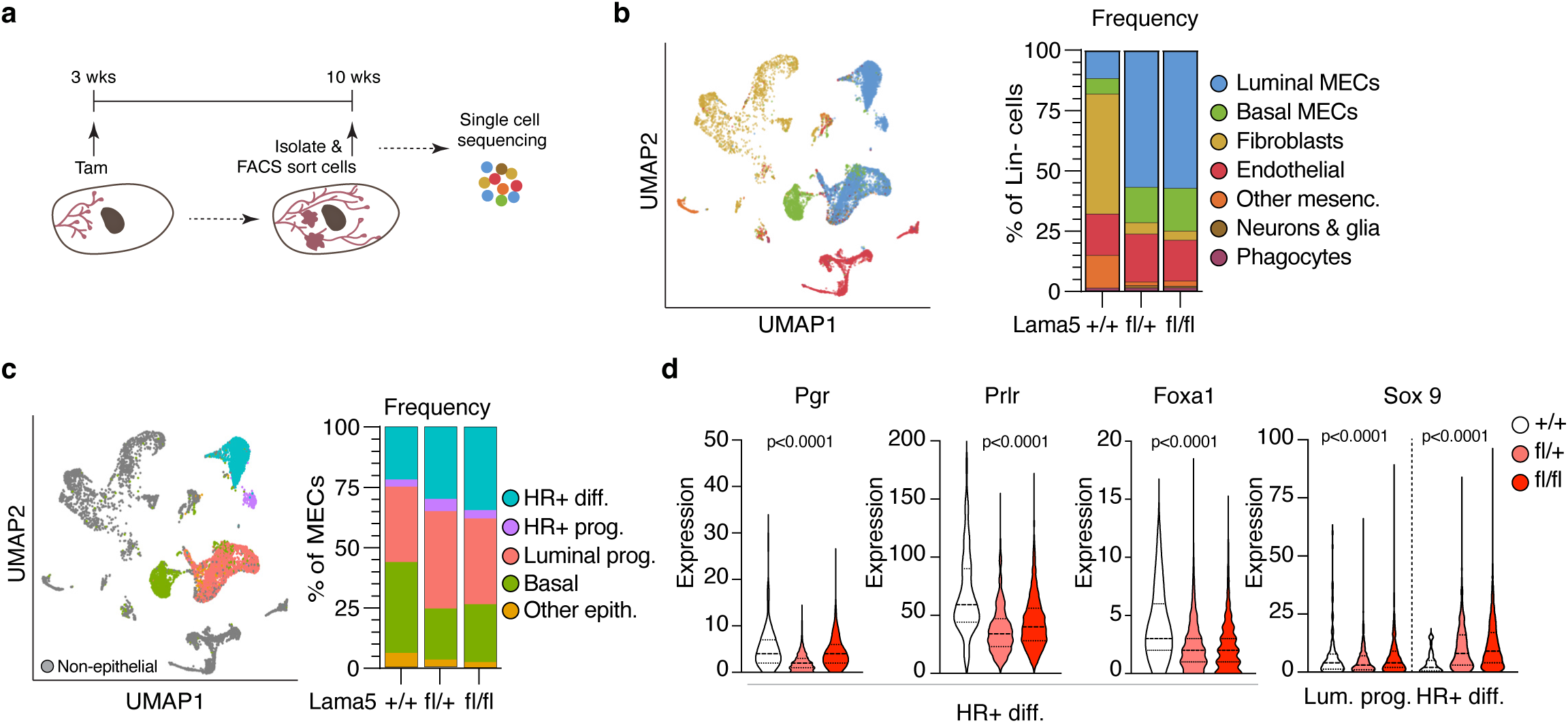
LAMA5 deletion alters epithelial cell populations. **a)** Schematic of the experimental set-up of single cell RNA sequencing from the MMTV-PyMT;K8-CreERT2;Lama5-flox mammary glands. **b)** UMAP showing clustering of MMTV-PyMT samples and identified clusters. In total ∼12650 cells analyzed. Graph showing frequency of the clusters identified in each genotype. **c)** Annotation of the cells to epithelial lineages according markers previously defined by *Bach et al. 2017*.Graph showing frequency of the lineages in each genotype. **d**) Violin plots showing expression of mature luminal markers Pgr, Prlr and Foxa1 in HR+ differentiated (HR+ Diff) population and Sox9 progenitor marker in luminal progenitor (Lum. prog.) and HR+ differentiated (HR+ Diff.) cell populations in indicated samples. Statistical analysis is performed using Kruskal-Wallis tests.

After dimension reduction, clustering, and cell type identification, we first explored the cellular composition of the samples by comparing the groups including all genotypes. The major clusters identified were luminal and basal epithelial cells, fibroblasts, endothelial cells, neurons and glia, and phagocytes (**Fig 4b, S4b**). The T;Lm5+/+ samples contained a drastically higher portion of fibroblasts and consequently less luminal and basal epithelial cells when compared to Lama5 targeted samples (**Fig 4b**). This possibly reflects the delay in tumour initiation in the T;Lm5fl/+ and T;Lm5fl/fl samples, as fibroblasts commonly constitute tumour stroma^34^ or differences in sampling. The changes in the cell types were however consistent across biological replicates with very similar frequencies in T;Lm5fl/+ and T;Lm5fl/fl samples (**Fig S4c**). Next, we annotated epithelial subtypes based on markers of MEC differentiation during pubertal gland growth^35^. Interestingly, we observed an increase in the luminal compartment, including frequency of HR+ differentiated, HR+ progenitor cells and luminal progenitor cells in both T;Lm5fl/+ and T;Lm5fl/fl samples (**Fig 4c; Fig S5**). Correspondingly, we observed in T;Lm5fl/+ and T;Lm5fl/fl samples a decrease in the frequency of basal epithelial cells (from 0,37 in T;Lm5+/+ to 0,21 in T;Lm5fl/+ and to 0,24 in T;Lm5fl/fl; **Fig 4c**). However, our immunofluorescence analyses of Krt8 and Krt14 in T;Lm5 tumours showed no significant alterations of luminal and basal cells between the genotypes (**Fig 3g,S3b**). This mismatch can arise either from methodological differences of measuring RNA and protein or from stricter definition of basal cell cluster in the sc-RNAseq analysis^35^.

Prompted by the observation of increased luminal populations, we additionally explored expression markers of mature mammary epithelial and progenitor cells. Interestingly, in Lama5 deficient tumours, cells classified as HR+ differentiated cells had lower expression of a subset of mature luminal markers, including *prolactin receptor* (*Prlr*) and *forkhead box A1* (*Foxa1*), a transcription factor that regulates differentiation of ER-positive luminal epithelial cells (**Fig 4d**)^35^. Furthermore, *progesterone receptor* (*Pgr*) expression was decreased specifically in T;Lm5fl/+ cells, indicating genotype-specific differences in the regulation of luminal cell states. Additionally, we observed an upregulation of progenitor markers *Sox9* ^35–38^ in HR+ epithelial cells from both T;Lm5fl/+ and T;Lm5fl/fl genotypes, whereas this increase was not detected in luminal progenitor cells (**Fig 4d**). Altogether, findings indicate that, in addition to the relative expansion of the luminal progenitor population, differentiated luminal epithelial cells exhibit fewer mature luminal features and increased progenitor-like characteristics, suggesting an overall shift toward a progenitor phenotype following Lama5 loss.

### Lama5 loss leads to changes in gene expression associated with FGF signalling

To further investigate whether the alterations in the luminal populations between *Lama5*-deficient and control tumours are linked to biological functions, we performed a single sample gene set enrichment analysis (ssGSEA) of the RNA sequencing data and compared the enrichment results in luminal epithelial cells by performing pathway clustering and similarity comparison to identify differentially enriched pathway clusters between the samples. In total, we identified 102 pathway clusters enriched in the T;Lm5 samples (**Fig 5a, FigS6a,b**). The major pathway cluster enriched in T;Lm5fl/+ and T;Lm5fl/fl samples in comparison to T;Lm5+/+ tumors was related to Fibroblast growth factor (Fgf) signalling pathways (**Fig 5a**) (*Fgfr1* ligand binding and activation, *Fgfr2* ligand binding and activation, and others [**Table S2]**). This was intriguing, as Fgf signalling has a significant role in mammary gland development and in breast tumorigenesis ^39^, and aberrant activation of Fgf receptor (Fgfr) via gene amplifications and mutations is a common feature of diverse tumours ^40^. Further analysis revealed that the fraction of luminal epithelial cells expressing *Fgf receptor 2* (*Fgfr2*) was increased particularly in T;Lm5fl/fl samples (**Fig 5b**). Taken together, our pathway clustering analysis reveals alterations in the inter-cellular communications in MMTV-PyMT tumours following early *Lama5* deletion.

**Figure 5.**
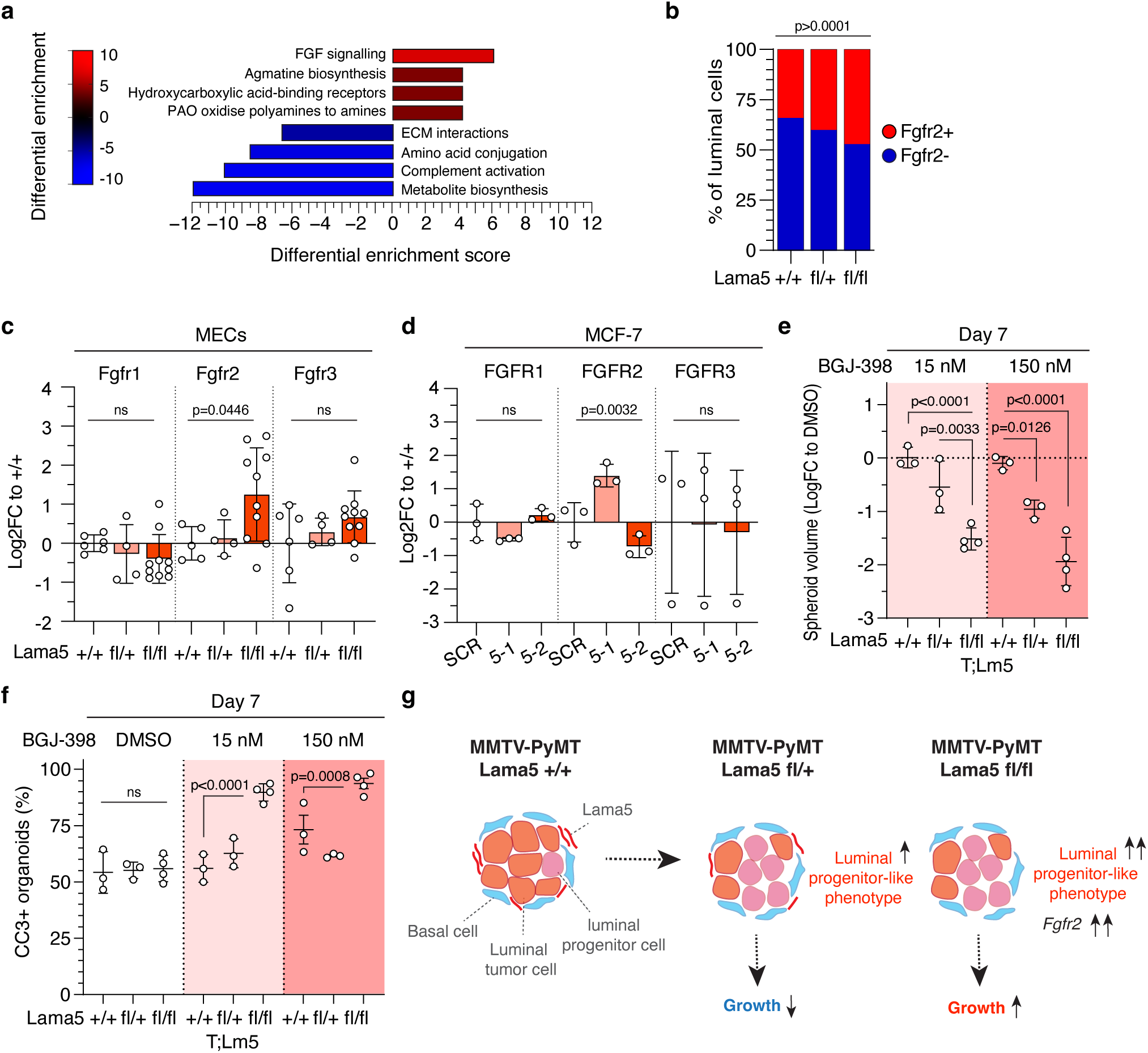
Inhibition of FGFR signaling reduces the growth of Lama5-deficient organoids. **a**) Differential pathway cluster enrichment of Lama5 fl/fl and Lama5 fl/+ tumors compared to Lama5 +/+ control tumors showing the top 4 pathways enriched in Lama5 deficient and control tumors. **b)** Frequencies of Fgfr2+ expressing luminal epithelial cells in each genotype. Statistical analyses perfomed using Chi Square test. **c)** qPCR analysis of *Fgfr1-3* in MECs of indicated genotypes. Each dot represents cells isolated from 1 mouse. Statistical analyses are performed using one-way ANOVA. **d)** qPCR analysis of FGFR1-3 in MCF-7 cells with LAMA5 downregulation (5-1, 5-2) or shCtrl (SCR). Each dot represents a biological replicate. Statistical analysis is performed using one-way ANOVA. **e)** Quantitation of organoid growth at day 7 in the indicated FGFRi treatments as log fold change of spheroid volume to DMSO. Each spot represents organoids isolated from 1 individual mouse. Statistical analysis is performed using two-way ANOVA and Tukey’s tests. **f**) Quantitation of caspase-3 (CC3) positive organoids at day 7 in the indicated treatments. Each dot represents organoids isolated from 1 mouse. Statistical analysis is performed using two-way ANOVA and Tukey’s tests. **g**) Schematic of the growth trajectory of the MMTV-PyMT mammary tumors after *Lama5* deletion.

### Inhibition of Fgf signalling reduces growth specifically in *Lama5*-deficient MECs

The enrichment of Fgf signalling in *Lama5*-deficient T;Lm5 samples led us to hypothesize the alterations in Fgf signalling may increase growth capacity especially in the tumors with biallelic loss of *Lama5*. To experimentally address this, we injected tamoxifen into T;Lm5fl/fl, T;Lm5fl/+ mice and T;Lm5+/+ control mice at the age of 8 weeks, corresponding to our earlier experiment where clear differences between T;Lm5fl/+ and T;Lm5fl/fl emerged in proliferation and carcinoma progression (**Fig 3c-d**), and isolated MECs 1 week later. We first investigated whether the upregulation of *Fgfr2* was induced following the acute postpubertal *Lama5* deletion in the mammary epithelium and performed qPCR of *Fgfr2,* along with *Fgfr1 and Fgfr3,* from the isolated cells. We observed a clear increase in the *Fgfr2* specifically in the T;Lm5fl/fl cells compared to T;Lm5+/+ and T;Lm5fl/+ cells (**Fig 5c**, **Fig S7a**). In line, FGFR2, but not *FGFR1* or *FGFR3,* expression was increased in the MCF-7 cell lines upon effective shRNA mediated downregulation of *LAMA5* (shLAMA5-1 **Fig 5d, Fig S7b**). However, shRNA with lesser knockdown efficiency did not induce a change (shLAMA5-2, **Fig 5d**). To further investigate whether the Fgfr pathway contributes to the growth properties the *Lama5* deficient epithelial cells, we grew the isolated MECs in 3D organoid culture for 7 days with or without Fgfr inhibitor infigratinib (BGJ-398 ^41^). In the control cultures the average diameter of the untreated T;Lm5fl/+ organoids was 32% smaller in comparison to T;Lm5+/+, yet the average T;Lm5fl/fl diameter was 20% larger in comparison to T;Lm5+/+ control (**Fig S7c-d**), indicating that the organoid culture can reproduce the growth phenotype seen *in vivo*. Intriguingly, while organoids from control mice remained unaltered, the size of T;Lm5fl/fl organoids was decreased with 15nM BGJ-398, and both T;Lm5fl/fl and T;Lm5fl/+ showed a clear size decrease with 150nM BGJ-398 concentration (**Fig 5e**). Moreover, the percentage of organoids with cells positive for active caspase-3 was increased in T;Lm5fl/fl organoids, whereas in T;Lm5fl/+ it remained similar to control cells (**Fig 5f**). While there was an increase in the frequency of apoptosis also in the 150nM BGJ-398 treated control cells, the number of apoptotic organoids was smaller compared to the T;Lm5fl/fl (**Fig 5f**). This suggests *Fgfr2* upregulation to present an acute and timepoint-independent consequence of *Lama5* loss, and that the growth-inhibitory and apoptosis-inducing effect of infigratinib, which selectively inhibits FGFR1, 2 and 3 ^41^, occurs likely via inhibition of Fgfr2. Finally, we determined whether upregulation of *Fgfr2* (**Fig 5c**) also occurs in the adenocarcinomas upon *Lama5* deletion (**Fig 3a)**. We found no differences in *Fgfr2* expression between the *Lama5* genotypes (**Fig S7e**), suggesting that *Fgfr2* is acutely upregulated in response to *Lama5* loss only in the early steps of luminal tumorigenesis. Taken together, our data shows that Fgf signalling, likely via the Fgfr2 receptor, is activated acutely in *Lama5*-lacking tumours and targeting Fgfr can specifically inhibit the growth of Lama5-deficient tumour cells, especially cells with biallelic *Lama5* deletion. Overall, this suggests altered Fgf signalling to represent a *Lama5*-independent compensatory mechanism that is utilized especially by tumours with biallelic *Lama5* loss, emphasising the interplay between ECM and epithelium during tumour progression.

## DISCUSSION

ECM has an essential role in cancer development and alterations in ECM composition and remodelling are known to occur during cancer progression ^2,4,42^. Here we demonstrate that BM component LAMA5 is overexpressed in human luminal breast cancers and its downregulation has a complex impact on tumorigenesis. Loss of *Lama5* strongly inhibits initiation of luminal mammary tumours *in vivo*. However, while reduced luminal *Lama5* expression slows tumour initiation, tumours with biallelic loss re-acquire normal tumour growth suggesting that they acquire other LAMA5-independent growth mechanisms (**Fig 5g**). Using sc-RNA sequencing we further show that *Lama5* deletion leads to a phenotype shift from differentiated luminal phenotype towards a luminal progenitor phenotype without luminal-basal conversion, along with changes in the expression of genes associated with Fgf signalling. Specifically, we identify that *Fgfr2* expression is upregulated in the *Lama5*-deficient luminal cells and inhibition of Fgfr signalling reduces growth and increases apoptosis in tumour cells with biallelic *Lama5* deletions.

We have previously demonstrated that *Lama5* is produced by HR+ luminal epithelial cells, and it is essential for HR+ luminal epithelial cell differentiation, function, and the growth of mouse mammary epithelium *in vivo* ^8^. Here we show that *Lama5* deletion in mammary tumour tissue decreases mature luminal markers *Prlr* and *Foxa1* and increases expression of progenitor markers *Sox9*. Deletion of luminal differentiation factors in mammary tumour models have previously been demonstrated to result in formation of tumours with basal phenotype and transcriptional programs that exhibit enhanced tumour development and shortened latency^43^. In contrast to these reports, we do not observe a switch from luminal to basal tumour type in the *Lama5*-deficient mice as measured by decrease in luminal *Krt8* and concomitant increase in basal *Krt14* positivity but rather see an increase in the luminal phenotype across the MECs. Emergence of double positive Krt8/Krt14 cells as an intermediate or hybrid stage between the two lineages prior to conversion to a basal-like phenotype has also been suggested ^29,32,44^. However, we did not detect a major shift in the amount of basal or double-positive epithelial cells, implying that loss of luminal differentiation factors does not directly result in transition to basal-like tumours. Similar results have been reported by deletion of other factors essential for luminal epithelial differentiation, such as Gata3, which also results in an increase in the luminal progenitor population and a decrease in tumour initiation capability, neither without apparent change towards basal-like histology^45^. However, luminal progenitor cell function appears important also for formation of basal-like tumours, as it was recently shown that deletion of the laminin-binding α6 integrin inhibits the growth of Brca1 and p53 deficient basal-like tumours by restricting expansion of luminal progenitor cells ^46^. Previous work on Brca1 tumours suggests that basal-like tumours arise from the luminal progenitor compartment ^30,47,48^ indicating that luminal progenitors function as the cell of origin or as a transitional population for basal tumours. Also, Sox9-positive luminal cells, which likely exhibit progenitor activity, have been shown to serve as cell of origin in LATS1/2 deficient basal-like tumours ^49^. Thus, it is possible that while the *Lama5*-deficient, luminal progenitor-rich tumours do not exhibit extensive basal gene expression they represent an intermediate state between basal and luminal in tumour development.

We show that deletion of *Lama5* leads to widespread deregulation of upregulation of Fgf signalling in luminal epithelial cells. Interestingly, we observed that this occurs acutely after *Lama5* deletion in the early tumors. Fgfr signalling has an essential role in mammary morphogenesis and in breast tumorigenesis by inducing growth and stem cell activity ^39,50,51^ and Fgfr overexpression has previously been linked to triple negative subtype, endocrine therapy resistance and mammary tumour progression ^51–53^. We demonstrate that inhibition of Fgfr activity in the *Lama5*-deficient tumour cells reduces their growth and induces apoptosis in biallelic-*Lama5*-deleted cells, suggesting that Fgf signalling can provide a surprising compensatory mechanism for the tumour promoting role of ECM remodelling. Whether upregulation of the Fgf signalling is a direct effect of *Lama5* loss or secondary to the alterations in the *Lama5*-deficient tumour ECM is unclear. Fgfs are bound to various ECM components, including heparan sulfate proteoglycans, fibronectin, and different laminin isoforms including LAMA5, which together function as a reservoir for the growth factors ^54–56^. Thus, modifications in the ECM organization, degradation or in laminin composition could alter the Fgf availability to the cells or the Fgf-Fgfr interactions ^56^ during tumorigenesis, triggering their deregulation. Hence, the outcome of Fgf signalling can be modified by the tumour ECM microenvironment. It is also possible that decrease in differentiated luminal and increase in progenitor gene expression is in part due to the increased Fgf signalling. For instance, Fgf signalling has been reported to activate the expression of Sox10, a close relative of Sox-9 ^57^, which both function as transcription factors in luminal progenitors ^58^. Therefore, we postulate that the amount of LAMA5 in the tumour microenvironment tunes tumour cell identity and signalling to an outcome favouring increased or decreased tumour growth (**Fig 5g**).

In line with its role as a tissue barrier, BM components have been considered to inhibit cancer development ^1,4^. However, these data and others recent results demonstrate that BM laminins can also be cancer promoting ^11,13–15^. We propose that certain components such as Laminin α5 are specifically important during tumour initiation, where they could participate in formation of a local BM microenvironment enabling the growth of hyperplastic cells. Spatially specialized BM microstructures have been observed for instance in developing dermal papilla and hair follicles and in Drosophila guiding embryonic tissue and imaginal disc development ^59–61^. The local ECM microenvironment has additionally been shown to regulate migration of normal and malignant cells ^62,63^. A systematic molecular profiling of the expression and localization of ECM during early tumorigenesis is needed to understand how distinct and local ECM niches are generated and how they contribute to cancer initiation. Our results underline that tumour microenvironment is a dynamic system and that decrease in one component can lead to global alterations in the tumour matrisome as well as in the inter-cellular communication. Finally, changes in tumour ECM can have a profound effect on tumour type and in the tumour cell populations that fuel the adaptation and progression of tumours.

## MATERIALS AND METHODS

### Experimental animals

#### Transgenic mice

All mice were of C57BL6/RccHsd background. MMTV-PyMT, Lama5 flox and K8-CreERT2 mouse lines have been previously described, and genotyping was performed using previously described primers ^23,24^. For K8-CreERT2 induction, a single dose of 2,5 mg tamoxifen in corn oil (20 mg/ml, both from Sigma) was injected intraperitoneally in 3-week-old mice and 5 mg tamoxifen in 8-week-old mice. Hyperplasia samples were acquired by collecting #4 mammary glands either from 10-week-old or 16-week-old mice. Adenocarcinoma samples were collected by excising all visible tumours from all mammary glands, when the mouse had developed at least one palpable tumour of 1 cm, which was the maximum tumor size permitted in and not exceeded in the experiments.

#### Fat pad transplantation

For the fat pad reconstitution assay mammary epithelial cells were isolated from 6-8 weeks old MMTV-PyMT donor mice as described below. 3-week-old female mice were anesthetized using 2-2,5% Isoflurane (Baxter) and the anterior part of #4 mammary gland, lymph node and bridge to #5 gland were surgically removed. To the remaining fat pad 10^5^ cells/gland were injected in 10 μl volume using Hamilton syringe. The wound was sutured using wound clips (Autoclip Physicians kit, Becton Dickinson), which were removed 1 week after the operation. The transplanted mice were sacrificed, and #4 mammary glands collected 6 weeks after transplantation when the recipient mice were 10 weeks old.

### Tissue histochemistry

#### Wholemounts

For wholemount staining, #4 inguinal mammary glands were fixed in 4 % PFA overnight and then stained for 48 hours in carmine-alumn staining solution (2% w/v Carmine, Sigma; 5 % w/v Aluminium Potassium Sulfate, Sigma). After the desired colour had developed, glands were mounted between glass, dried for 1-2 weeks, and imaged with Leica S9i stereo microscope with 1.5-2x magnification.

#### H&E stainings

#4 mammary glands and adenocarcinomas were fixed in 4 % PFA overnight, embedded in paraffin, cut into 5 µm sections, deparaffinized, and rehydrated prior to staining. H&E stainings were performed in the Finnish Center for Laboratory Animal Pathology (FCLAP) at the University of Helsinki. Imaging of H&E-stained sections was performed using a Pannoramic 250 Flash II high-throughput brightfield scanner (3DHISTECH) with a 20X (NA 0.8) objective and Pannoramic Viewer software.

#### Immunofluorescence stainings

For immunofluorescent stainings, antigen retrieval was performed by placing the sections in a microwave in warm 1 mM EDTA for 8 minutes and then incubated at room temperature for 20 minutes. Unspecific antibody binding sites were blocked with a blocking buffer (10% normal goat serum [NGS, Gibco] in PBS) for 2 hours at room temperature. Primary antibodies (anti-cytokeratin-8 [TROMA-1, Developmental Studies Hybridoma Bank], anti-cytokeratin-14 [PRB-155P, BioLegend] and Ki-67 [ab16667, Abcam]) were diluted in the blocking buffer 1:300 and incubated on the sections overnight at 4°C. The following day, secondary antibodies (Goat anti-Rat, Alexa Fluor™ 633 and Goat anti-Rabbit, Alexa Fluor™ 488 [Thermo Fisher Scientific]) diluted in the blocking buffer 1:500 were incubated on the samples for 2 hours at room temperature light protected, followed by 1 μg/ml Hoechst (Sigma-Aldrich, H33342) nuclear counterstaining for 20 minutes at room temperature, light protected. Between all steps, the sections were washed with PBS for 15 minutes. The sections were mounted on glass using Immu-Mount (Thermo Fisher Scientific) and imaged at the Genome Biology Unit (GBU) at the University of Helsinki using 3DHISTECH Pannoramic 250 FLASH II digital slide scanner with 40X/0.95 NA objective.

#### RNA in situ hybridization

RNAscope 2.5 HD Reagent Kit-Brown or an RNAscope Multiplex Kit (Advanced Cell Diagnostics Srl) was used according to manufacturer’s instructions to perform RNA in situ hybridization on human breast cancer tissue microarray sections (Biomax BC081116d) and mouse mammary tumour sections. The sections were pretreated with RNAscope Target Retrieval Reagents, dried overnight, and used the following day. Probes for either human laminin α5 or mouse laminin α1, α3, α4 or α5, (Catalog #530691, #494931, #494921, #494901 and #494911, Advanced Cell Diagnostics) were hybridized for 2 h at 40°C. Thereafter, signal amplification hybridization was performed, followed by detection with DAB (3,3′-diaminobenzidine) and hematoxylin counterstaining. Samples were imaged using a Leica DM2000 LED equipped with a Leica MC 190 HD camera unit and a 40x/0.65 air objective. Both TMA and mouse mammary tumour sections were imaged with Pannoramic 250 Flash II high-throughput brightfield scanner (3DHISTECH) with a 20X (NA 0.8) objective and Pannoramic Viewer software.

### Culture and analysis of cell lines

#### Cell culture

MDA-MB-231 cells were grown in Dulbecco modified Eagle’s medium (DMEM, Sigma-Aldrich) containing 10% of fetal bovine serum (FBS, Gibco), 2 mM glutamine (Sigma-Aldrich), 100 U/ml penicillin (Orion) and 100 μg/ml streptomycin (Sigma-Aldrich). MCF-7 cells were grown in the same media as described for MDA-MB-231, except with additional 5 μg/ml human insulin (Sigma-Aldrich) and 1 mM sodium pyruvate (Sigma-Aldrich). Both cell lines were grown at a humidified 5% CO^2^ 37°C incubator, and passaging was performed after >80% confluency.

Single cell suspensions were prepared from MCF-7 and MDA-MB-231 cells by incubating the cells in 0.05% Trypsin-EDTA (Difco, J.T. Baker) at 37°C for 3-5 minutes, followed by centrifugation at 300 G for 3 minutes. The pellets were resuspended with the appropriate culture medium and filtered through a 40 μm strainer (Fisher Scientific). MCF-7 cells were additionally pushed through a 30 G needle (Becton, Dickinson).

### Lentivirus production and transduction

293FT cells cultured in antibiotic-free media (DMEM, 10% FBS, 2 mM glutamine) were transfected in 5% Lipofectamine™ 2000 (Invitrogen) using packaging plasmids Δ8.9, CMV-VSVg, and a transfer plasmid (pLKO.1 with non-targeting shScramble SHC002 [Sigma-Aldrich] or a *LAMA5*-targeting shLAMA5-1 or shLAMA5-2 shRNA construct, or LentiCRISPRv2 gSCRA or pLentiCRISPRv2 gLAMA5) followed by 72-hour incubation. Lentivirus was collected from the media through 0.45 μm PES filters (Whatman). MCF-7 and MDA-MB-231 cells were transduced in the appropriate cell culture medium supplemented with 10 mg/ml Polybrene transfection agent (Sigma-Aldrich, H9268) for 48 hours, followed by two-week selection in media supplemented with 2 mg/ml puromycin (Sigma-Aldrich). Primary MMECs were transduced on low-adhesion culture plates for 72 hours before transplantation or placing into organoid culture. Downregulation was validated with either LAMA5 protein immunofluorescence staining of cells grown overnight in 3D suspension cultures (cell lines), as described below, or with qPCR (primary MMECs).

### Live cell imaging

25000 MCF-7 or 75000 MDA-MB-231 single cells were plated onto well plates, incubated overnight, and then placed into a Cell-IQ® live imaging system (CM Technologies) for 72 hours. Each well was imaged from three positions with equal cell distribution once every 30 minutes using a built-in Qimaging Retiga EXi camera with 10x/0.30 objective.

### EdU incorporation assay

50000 MCF-7 single cells were plated on 15 mm glass coverslips in the appropriate cell culture medium and incubated for 24 hours. The medium was then changed to a fresh medium supplemented with 10 μM 5-Ethynyl-2’-deoxyuridine (EdU) for 2 hours and fixed with 4% PFA for 10 minutes at room temperature.

### 3D suspension culture

4000 single cells suspended into 1 ml of 1% w/v solution of methyl cellulose (Sigma-Aldrich) in the appropriate cell culture medium were plated onto each well of 24-well ultra-low-adhesion cell culture plates (Corning). The growing spheroids were inspected on the following day from the plating to ensure equal plating confluency and thereafter cultured for 1 or 10 days. The centre position of each well was imaged using a Nikon Eclipse TS100-F trinocular inverted microscope equipped with a Nikon Digital Sight DS-L3 camera control unit, with 4x/0.13 air objective. The spheroids were fixed by diluting the methyl cellulose suspension 1:1 with the appropriate cell culture medium, centrifuging at 500 G for 5 minutes, and resuspending the pellets into 4% PFA for 25 minutes in room temperature. Post-fixation, the spheroids were centrifuged again at 500 G for 5 minutes and resuspended in 0.2 % BSA in Dulbecco’s PBS.

### Cell isolation

Mouse mammary epithelial cells (MMECs) were isolated from mammary glands of mice with or without tumour growth. Shortly, the glands #3-#5 were dissected, and the lymph node in #4 gland removed. The tissues were mechanically chopped and incubated with 0.01 mg of Collagenase A (Sigma) per 1 g of tissue in Advanced DMEM/F12 growth media (Life Technologies) containing 2,5% FCS, 5 μg/ml insulin, 50 μg/ml gentamicin and 2 mM glutamine in gentle shaking (120 rpm in environmental shaker) at 37 °C for 2-2,5 hours. The resulting cell suspension was then first centrifuged 400 RCF for 10 minutes and consecutively pulse centrifuged 3-5 times 400 RCF to get a preparation free of cells other than MMEC organoids. Next organoids were trypsinized with 0.05% Trypsin-EDTA (DIFCO, J.T. BAKER) for 5-10 minutes and drained through 70 μm cell strainer to obtain single cells and centrifuge at 300 RCF for 5 minutes (Becton Dickinson). Cell pellet was resuspended in MMEC growth media (Advanced DMEM/F12 media containing 5 μg/ml insulin, 1 μg/ml hydrocortisone, 10 ng/ml mouse EGF, 2 mM glutamine, 50 μg/ml gentamycin and penicillin and streptomycin (all from Sigma) supplemented with 10% FCS (Gibco) or in 0,2 % BSA in Dulbecco’s PBS for FACS sorting.

### 3D cell culture of primary and tumour MMECs

Isolated MMECs were cultured overnight on 24-well ultralow adhesion plates (Corning Costar) prior to 3D cultures. 3D organotypic culture was performed in growth factor reduced basement membrane from Engelbreth-Holm-Swarm (EHS) mouse sarcoma (Matrigel^tm^, Becton Dickinson), which was prepared according to manufacturer’s instructions. MMECs were collected from ultralow adhesion plates, centrifuged 400 RCF for 3 minutes and suspended with liquid Matrigel and plated onto 8-chamberslides in Matrigel mixture drops approximately 2000 cells/drop. Organoids were grown in MMECs growth media, which was refreshed every 3-4 days. For Fgfr inhibition experiments, 15μM or 150μM Infigratinib (BGJ398, Selleckchem) or DMSO as carrier control was added to the cells on day 1 after seeding.

### qPCR and analysis

#### RNA isolation and qPCR

RNA was isolated from tissues or cells using NucleoSpin™ RNA isolation kit (Macherey-Nagel) with DNase treatment, according to the manufacturer’s instructions. cDNA was synthesized from 500 ng of RNA using Revert Aid cDNA synthesis kit (Thermo Fisher Scientific) with Random Hexamer primers according to the manufacturer’s instructions, in a T100™ Thermal Cycler (Bio-Rad).

RT-qPCR reaction was performed in a CFX384 Touch Real-Time PCR Detection System (Bio-Rad) in Power SYBR™ Green PCR Master Mix (Thermo Fisher Scientific) according to the manufacturer’s instructions. Primer sequences are presented in the Table 1 below. RT-qPCR results were quantified with Bio-Rad CFX Maestro 1.1 software, using GAPDH and ACTB as house-keeping genes. The gene expression results are presented as logarithmic fold changes (LogFC) compared to the control sample.

**Table 1.**
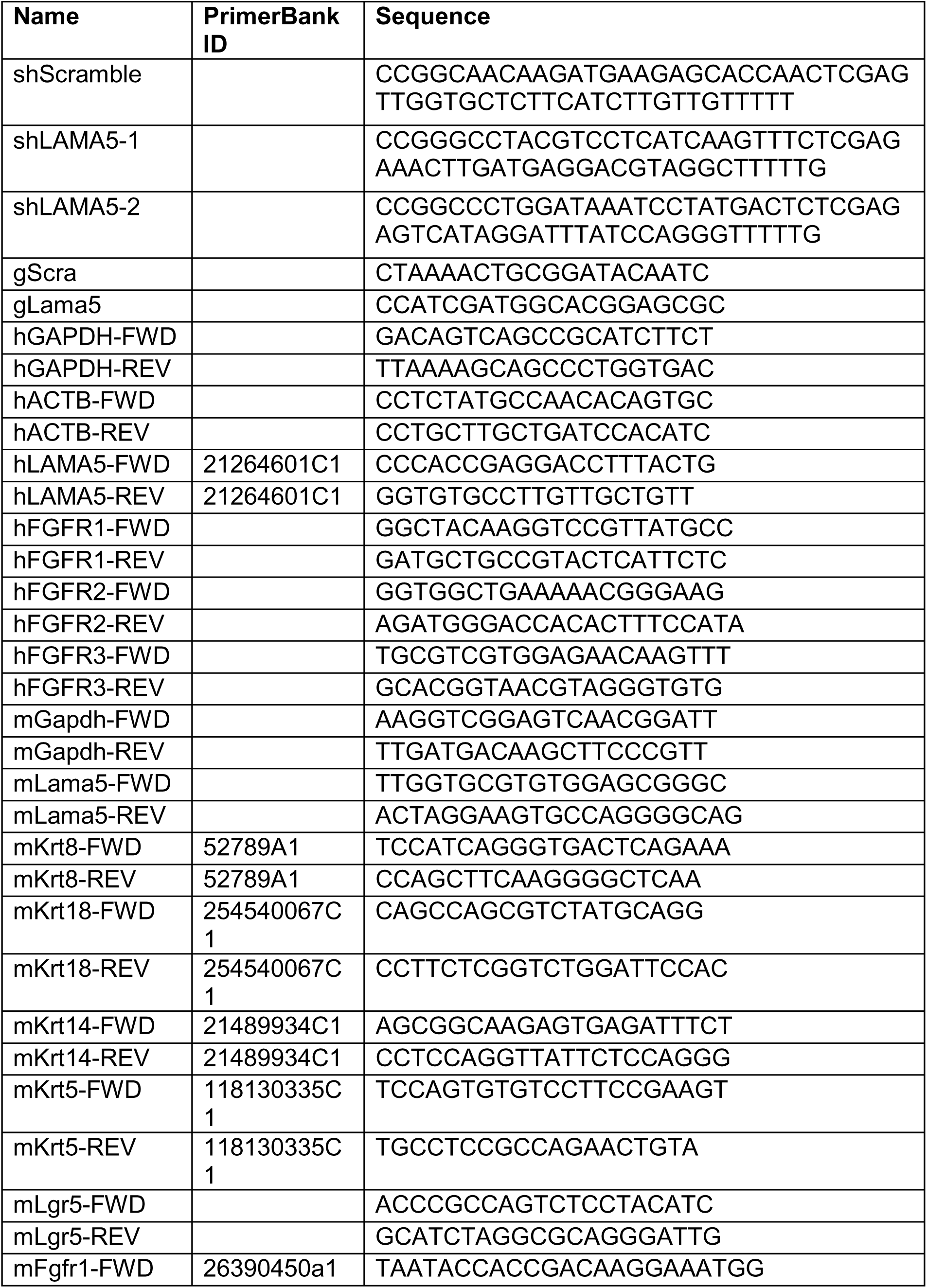
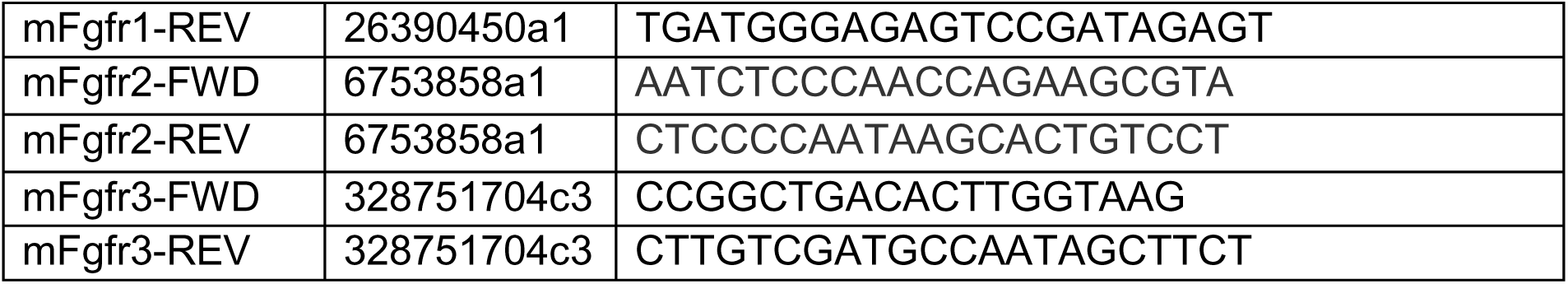

### Immunocytochemistry

#### Spheroid immunostainings

Unspecific binding sites were blocked with a blocking buffer of 10% NGS (Thermo Fisher Scientific) in 0.2% BSA in Dulbecco’s PBS for 90 minutes at room temperature. Primary antibody (anti-laminin-alpha-5 [4C7, Millipore]) diluted 1:500 in the blocking buffer was incubated overnight at 4°C. The following day, the spheroids were incubated with the secondary antibody (Goat anti-Mouse, Alexa Fluor™ 546 [Thermo Fisher Scientific]) diluted in the blocking buffer 1:500, for 2 hours at room temperature, light protected, and thereafter the nuclei were counterstained with 1 μg/ml Hoechst (Sigma-Aldrich, H33342) for 20 minutes at room temperature, light protected. All incubations and washes were done in gentle shaking, and reagents were changed by twice centrifuging the spheroids at 300 G for 3 minutes and resuspending the pellet with 0.2% BSA in Dulbecco’s PBS. The spheroids were mounted in 1:1 diluted Immu-Mount (Thermo Fisher Scientific) onto a MatTek 35 mm glass bottom dish (MatTek Slovakia) with a glass coverslip on top and imaged using a Leica TCS SP8 confocal microscope equipped with an HC PL APO CS2 20x/0.75 immersion oil objective and a Leica Hybrid Detector (HyD) photon counting mode.

#### Organoid immunostainings

3D organoids were fixed with 2% PFA for 20 minutes at room temperature and thereafter washed with PBS. Organoids were permeabilized with 0.25% Triton X-100 in PBS for 10 minutes at + 4°C and thereafter washed with PBS. The non-specific binding sites were blocked in immunofluorescence (IF) buffer (0.2% Triton-X, 0.05 % Tween, 0.1 % BSA in PBS) supplemented with 10% normal goat serum (Gibco) for 1-2 hours. The primary antibody (Active caspase-3, Cell Signaling Technologies) was incubated in the IF buffer overnight at +4°C. Following the incubation, structures were washed three times with IF buffer, 15 minutes each wash, and then incubated with secondary antibody (Goat-anti rabbit Alexa Fluor 488 [Thermo Fisher Scientific]) diluted in IF buffer 1:300 for 45 minutes at RT. The samples were washed with IF buffer and the nuclei were counterstained with Hoechst33342.

#### EdU incorporation staining

The cells were permeabilized with 0.1% v/v Triton X100 (Sigma-Aldrich) in PBS for 10 minutes and nuclei incorporating EdU were stained using Click-iT™ EdU Cell Proliferation Kit (Invitrogen, C10340) with Alexa Fluor™ 647 according to the manufacturer’s instruction, followed by nuclear counterstaining with 1 μg/ml Hoechst (Sigma-Aldrich, H33342) for 20 minutes at room temperature, light protected. Finally, the coverslips were mounted on a microscope slide using Immu-Mount (Thermo Fisher Scientific) and imaged using a Leica DM5000B-CS fluorescent microscope with a 20x/0.70 air UV objective.

### Image analysis and quantification

#### Wholemount quantification

Area of proximal tumour hyperplastic growth on the individual 10-week-old or 16-week-old glands as mm^2^ was quantified by first manually annotating tumour areas from the wholemount images in GIMP, and then selecting the annotated areas within the distance of 4.27 mm from the beginning of the ductal tree (approximately 1/5 of the median length of a #4 mammary gland) with the “Radial Area”-protocol in Tonga v0.1.9 ^19^. Glands with maximal ductal growth less than 4.27 mm from the beginning of the ductal tree were excluded. Ductal network lengths were quantified using GIMP, ImageJ, and a custom JavaScript-script. Fat pad invasion was defined as the distance from the beginning of the ductal tree to the farthest tip of the epithelium, along the gland morphology. Ductal growth density was calculated as a ratio of ductal network length to fat pad invasion length. Samples with a palpable tumour and/or proximal lesion area exceeding 2 mm^2^ (25^th^ percentile of average proximal lesion area in all 16-week-old T;Lm5 samples) were considered to have an “advanced lesion”.

#### Quantification of cell proliferation

The total number of adherent non-apoptotic nuclei and the proportion of nuclei with EdU incorporation were analysed automatically from the images using the “Count the ratio of positive nuclei”-protocol in Tonga v0.1.2 ^19^ with the intensity parameter set as 2%.

#### Quantification of live cell growth

Confluency values at each timepoint were measured automatically using the “Area size measurement” function in the Cell-IQ Analyzer software v4.4.0. Starting timepoints were adjusted to an equal starting confluency between all samples. The average cumulative growth was calculated for each sample and normalized as a ratio to the final average cumulative growth of the control sample at 48 hours to obtain relative growth values, which were used to fit growth curves. Doubling times were calculated by fitting unnormalized confluency values. Non-linear least squares regression with an exponential growth model was used for fitting.

#### Quantification of spheroid growth

The total image area occupied by the spheroids in each sample was analysed automatically using the “Organoids”-protocol in Tonga v0.1.9 ^19^ with the target size of 25 for MDA-MB-231 images, and 50 for MCF-7 images. Three-dimensional growth of spheroid volume was estimated from two-dimensional plane area by the logarithm base ^3^√4 transformation, and the data is presented as logarithmic fold changes (LogFC) compared to the control sample.

#### LAMA5 fluorescence intensity quantification

LAMA5 fluorescence intensity was quantified as channel sum or mean intensity from spheroid stack images at a circular area of the middlemost section for each spheroid manually in ImageJ v1.53.

#### Cytokeratin fluorescent intensity quantification

Hyperplasia or adenocarcinoma areas from each sample were manually annotated from the images with 3DHISTECH CaseViewer v2.4.0 and a pyramid level 3 for hyperplasia or level 4 for adenocarcinoma was extracted using the MiraXtractor software (https://github.com/avritchie/miraxtractor). Uneven illumination correction, background correction, and tissue area segmentation were performed using the “IF tissue intensities” protocol and “in Tonga v0.1.9 ^19^. For protocol parameters, combined non-tissue removal method and tissue scale factor of 40 for the hyperplasia and 20 for the adenocarcinoma images were chosen. The proportion of tissue area positive for a given channel was defined as the proportion of tissue exceeding 25% channel signal intensity.

#### Ki-67 quantification

Hyperplasia or adenocarcinoma areas were manually annotated from the images with 3DHISTECH CaseViewer v2.4.0. Using the “IF nuclear intensity measurements” protocol in Tonga v0.1.9 ^19^, the images were automatically pre-processed with uneven illumination and background correction, the nuclei segmented, and the signal intensities of the individual nuclei measured. Pyramid level 0 and nuclear target size of 40 were used as protocol parameters. Ki-67 positivity was defined for each sample as the proportion of tumour nuclei with nuclear sum intensity of the signal channel exceeding 25.

#### RNAScope quantification

RNA expression of laminin α chains was estimated from the RNAScope images by performing epithelial tissue area segmentation and relative chromogen intensity measurement by channel deconvolution with the “Chromogen signal intensity” protocol in Tonga v0.1.9 using 85 as the chromogen sensitivity and 85 as the segmentation sensitivity parameter for the default methods. The results are presented as mean chromogen intensity of the analysed tissue area as parts per thousand (‰). Mouse samples were further normalized between mice to arbitrary units (A.U.) using mean chromogen intensity of normal mammary epithelial tissue in each mouse.

#### Histological grading and classification of tumours

H&E-stained sections of tumours were analysed by two blind observers and classified as solid, tubulopapillary, tubular, cystic-papillary or cystic-cribriform subtypes according to previously described criteria^24^. Grade of the tumours was determined by combining the consensus score of the observers on tubule formation, nuclear pleomorphism, and the number of mitoses per 10 high power fields (Table S1 for details).

#### Active caspase-3 quantification

The frequency of organoids containing ≥1 active caspase-3 positive cells was manually quantified from the immunostained samples. 60-200 organoids per sample were counted in each experiment.

### Single cell RNA sequencing and data analysis

#### Collecting of MMECs for sc-RNA seq

Primary cells were isolated from mice with the following genotypes: K8;Lama5 fl/fl (2 mice), K8;Lama5 fl/+ (1 mouse) and K8;Lama5 +/+ (1 mouse), hereafter collectively *WT*, and T;Lm5 fl/fl (2 mice), T;Lm5 fl/+ (1 mouse) and T;Lm5 +/+ (2 mice), hereafter *PyMT*. Single cell suspension of isolated primary MMECs was resuspended in 0,2 % BSA in Dulbecco’s PBS and the cells were incubated with either of the following set of primary antibodies: CD31-BV421, CD45-BV421 and Ter119-BV421, (from Becton Dickinson). All antibodies were used 1:500 and incubated on ice for 30 minutes. Cells were washed with PBS and re-suspended in 0,2 % BSA in Dulbecco’s PBS with 2 μM Sytox Blue (LifeTechnologies, Thermo Fisher) to exclude dead cells. Sorting was performed with BD FACSAria Fusion Flow Cytometer (Laser 405nm, 488nm, 561nm, 633nm) (Beckton Dickinson). FlowJo V10 was used for post-analysis of sorted cells.

#### sc-RNA sequencing library preparation and data processing

Altogether 50 000 FACS sorted, lineage negative live cells per sample were processed for library preparation in 0,04% BSA in PBS. Before library preparation the cell counts, and viability was determined by Luna cell counter (Logos Biosystems). Single cell gene expression profiles were studied using 10x Genomics Chromium Single Cell 3’RNAseq platform. The Chromium Single Cell 3’RNAseq run and library preparation were done using the Chromium Next GEM Single Cell 3’ Gene Expression version 3.1 Dual Index chemistry. The Sample libraries were sequenced on Illumina NovaSeq 6000 system using read lengths: 28bp (Read 1), 10bp (i7 Index), 10bp (i5 Index) and 90bp (Read 2). Data processing was performed using 10x Genomics Cell Ranger v4.0. The pipeline “cellranger mkfastq” was used to produce FASTQ files and “cellranger count” to perform alignment, filtering and UMI counting. mkfastq was run using the Illumina bcl2fastq v2.2.0 and alignment was done against mouse genome mm10. “Cellranger aggr” pipeline was used to combine data from multiple samples into an experiment-wide gene-barcode matrix and analysis.

#### sc-RNA seq data analysis

All data were analysed in R using the Seurat package^64^ for most operations. Samples were filtered for low counts and contaminant mitochondrial genes, and cell identities determined by the supervised combination of automated cell deconvolution labels with SingleR^65^ and MEC signature^35^ enrichment from the Semi-supervised Category Identification and Assignment (SCINA) package^66^, with settings as follows: max_iter = 100, convergence_n = 10, convergence_rate = 0.999, sensitivity_cutoff = 0.9. Quality control yielded 10, 262 cells (wt) and 12, 654 cells (MMTV-PyMT) in total for analysis. When requested, epithelial cell subtyping was performed using the standard Seurat label transfer method with references from Bach *et al*^35^. Unsupervised dimension reduction and cluster analysis was performed with Uniform Manifold Approximation and Projection (UMAP) ^64^.

#### ssGSEA and pathway enrichment analysis

Functionally related enriched pathways were grouped together by calculating pathway similarity and performing walktrap-clustering with simplifyEnrichment R package version 1.12.0^67^ for all pathways found enriched through ssGSEA across the samples as well as for each sample individually. Differential pathway enrichment between samples was compared by calculating a cluster similarity matrix based on gene overlap for each pair of clusters and performing a diagonal matrix transformation to obtain a differential enrichment score for each cluster for all genotypes. The calculated scores were weighted by multiplying with cluster size and maximum cluster similarity across genotypes to obtain the final differential enrichment scores for each pathway cluster for each genotype in comparison to the other genotypes

#### TCGA expression analysis

RNA-sequencing and gene copy number data from the TCGA-BRCA project were downloaded from the GDC portal. A primary tumour from each case was classified into tumour subtypes using HR, PR, and HER2 IHC status of each sample from the clinical data. Cases with missing IHC status were excluded. For analysis by tumour stage, pathological stage by AJCC criteria was used and cases without staging information were further excluded. RNA-sequencing counts were normalized using DESeq2 1.38.3, and normalized counts were compared between the tumour subtypes or tumour stages using Kruskal-Wallis ANOVA tests.

## Quantification and statistical analysis

Data is presented as mean ± SD or SEM from at least three independent experiments, unless otherwise stated in the figure legend. All statistical tests were performed using GraphPad Prism 10.

## DECLARATIONS

### Ethics approval

All animal studies were approved by the National Animal Ethics Committee of Finland (ELLA; Licence # ESAVI-18179-2020) and conducted in the Laboratory animal centre (LAC) of the University of Helsinki under institutional guidelines.

### Availability of data and materials

The datasets used and analysed during the current study are available from the corresponding author on reasonable request. Sc-RNA sequencing files have been deposited to Figshare and can be accessed through the link: https://figshare.com/s/e490f9d61963efe4f207

## Author contributions

AR – investigation, methodology, formal analysis, visualization, writing – original draft, writing – review & editing

PD – investigation, methodology, formal analysis, visualization, writing – original draft, writing – review & editing

EK – investigation, methodology

JL – methodology

LB – resources

VI – resources, methodology

PK – supervision, funding acquisition, writing – review & editing

JE – investigation, formal analysis, visualization, supervision, funding acquisition, writing – original draft, writing – review & editing

## Acknowledgements

1. J. Bärlund and M. Simula are thanked for excellent technical assistance. We thank all the members of Katajisto and Englund laboratories for comments and discussion. Light Microscopy Unit (LMU) at the Institute of Biotechnology, University of Helsinki is acknowledged for assistance with microscopy. Finnish Center for Laboratory animal Pathology (FCLAP) is acknowledged for histology services. Genome biology Unit (GBU), supported by HiLIFE and the Faculty of Medicine, University of Helsinki, and Biocenter Finland is acknowledged for scanning of histological slides. The authors would like to thank FIMM Single-Cell Analytics unit supported by HiLIFE and Biocenter Finland for single-cell RNA sequencing services. Part of this study was carried out with the support of HiLIFE Laboratory Animal Center Core Facility, University of Helsinki. The results published here are in part based upon data generated by the TCGA Research Network: https://www.cancer.gov/tcga.

