## Supplemental Figure 1 for "Tumour-derived LAMA5 is critical for tumour initiation and controls progression and phenotype in luminal breast cancer"

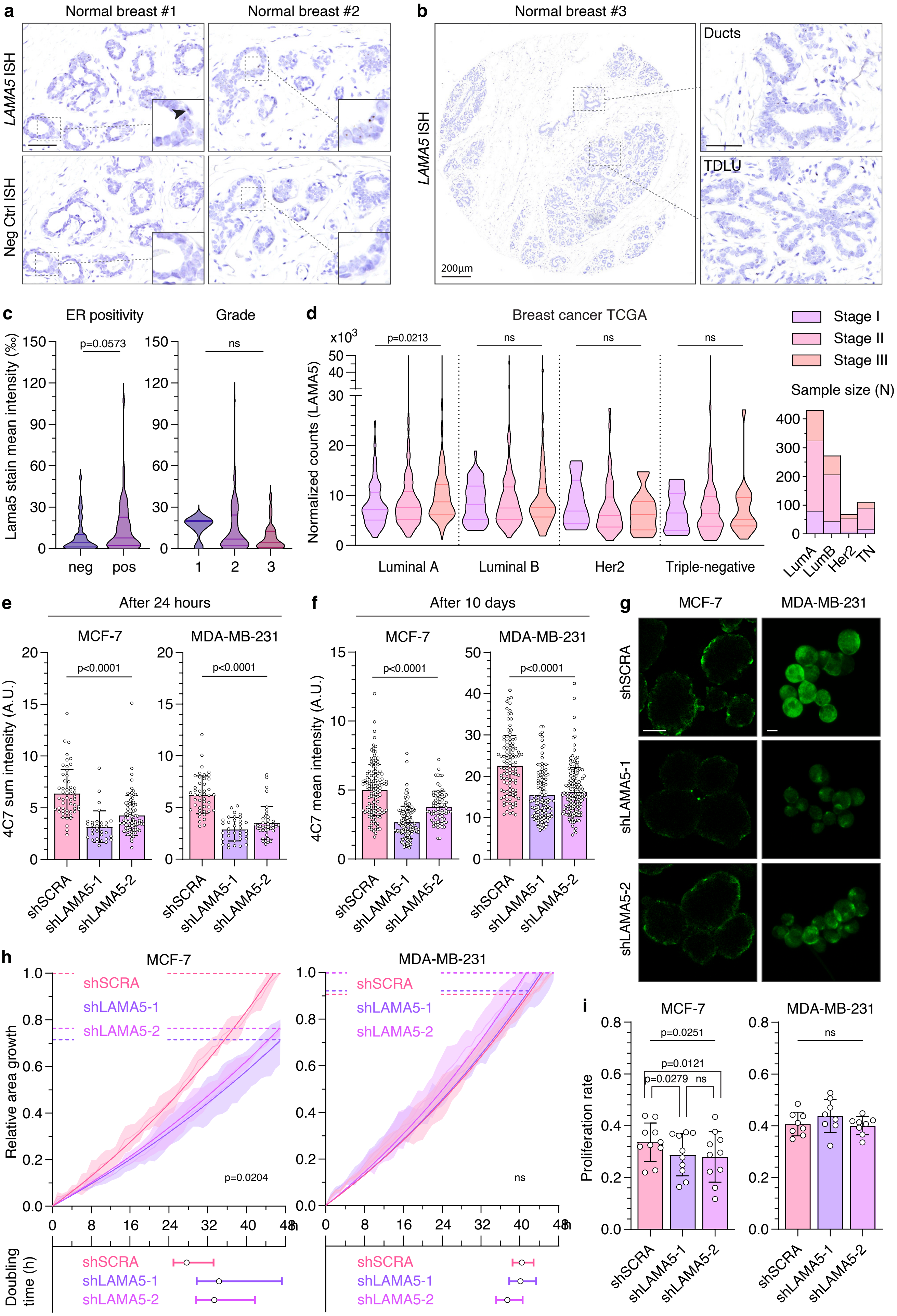

a
Normal breast #1
Normal breast #2
LAMA5 ISH
Neg Ctrl ISH
b
Normal breast #3
LAMA5 ISH
Ducts
TDLU
200μm
c
ER positivity
Grade
Lama5 stain mean intensity (%)
p=0.0573
ns
neg
pos
1
2
3
d
Breast cancer TCGA
Normalized counts (LAMA5)
x10^3
p=0.0213
ns
Luminal A
Luminal B
Her2
Triple-negative
Stage I
Stage II
Stage III
Sample size (N)
LumA
LumB
Her2
TN
e
After 24 hours
MCF-7
MDA-MB-231
4C7 sum intensity (A.U.)
p<0.0001
shSCRA
shLAMA5-1
shLAMA5-2
f
After 10 days
4C7 mean intensity (A.U.)
g
MCF-7
MDA-MB-231
shSCRA
shLAMA5-1
shLAMA5-2
h
MCF-7
MDA-MB-231
Relative area growth
p=0.0204
ns
Doubling time (h)
i
Proliferation rate
p=0.0251
p=0.0121
p=0.0279

**Supplemental figure 1. LAMA5 is overexpressed in human luminal breast tumours and required for growth of luminal breast cancer cells.**

**a)** In situ hybridization (ISH) of *LAMA5* and negative control probe in normal human breast. Scale bar 50 μm. **b)** Representative images of ISH of *LAMA5* TDLU vs duct. Scale bar 50 *μ*m. **c)** *LAMA5* positivity in TMA samples assigned to groups according to ER status (negative or positive; left graph) and grade (1-3; right graph). ER- n=35, ER+ n=76. Grade 1 n=5, 2 n=68, 3 n=25. Statistical analysis is performed using a Mann-Whitney test and a one-way ANOVA (Kruskal-Wallis test) d) Lama5 expression according to subtypes and tumour stage from The Cancer Genome Atlas. Kruskal-Wallis test was used for statistical tests. e-f) Downregulation of LAMA5 in MCF7 cells and in MDA-MB-231 cells shown using LAMA5 protein intensity measurements with 4C7 antibody after 24 hours (e) or 10 days (f) of low-adhesion culture. Each dot represents a single cell (e) or spheroid (f). g) Representative images of 4C7 IF of MCF7 and MDA-MB-231-cells expressing either LAMA5 shRNA or Scrambled shRNA grown as spheroids for 10 days in low-adhesion culture. Scale bar MFC-7 50 μm, MDA-MB-231 10 μm. h) growth curves for MCF-7 or MDA-MB-231 expressing either LAMA5 shRNA or Scrambled shRNA for 48 hours in monolayer culture, relative to Scrambled shRNA. Average growth curve, standard deviation, and fitted curves for with exponential growth equation are visible. Mean population doubling times in hours are visible in the bars below. RM one-way ANOVA on fitted growth rates was used for statistical testing. i) proliferation rate of MCF-7 or MDA-MB-231 cells expressing either LAMA5 shRNA or Scrambled shRNA after a 2-hour EdU pulse. Each dot represents a repeated experiment. RM one-way ANOVA and Fisher's LSD tests were used for statistical testing.
