## Supplemental Figure 2 for "Tumour-derived LAMA5 is critical for tumour initiation and controls progression and phenotype in luminal breast cancer"

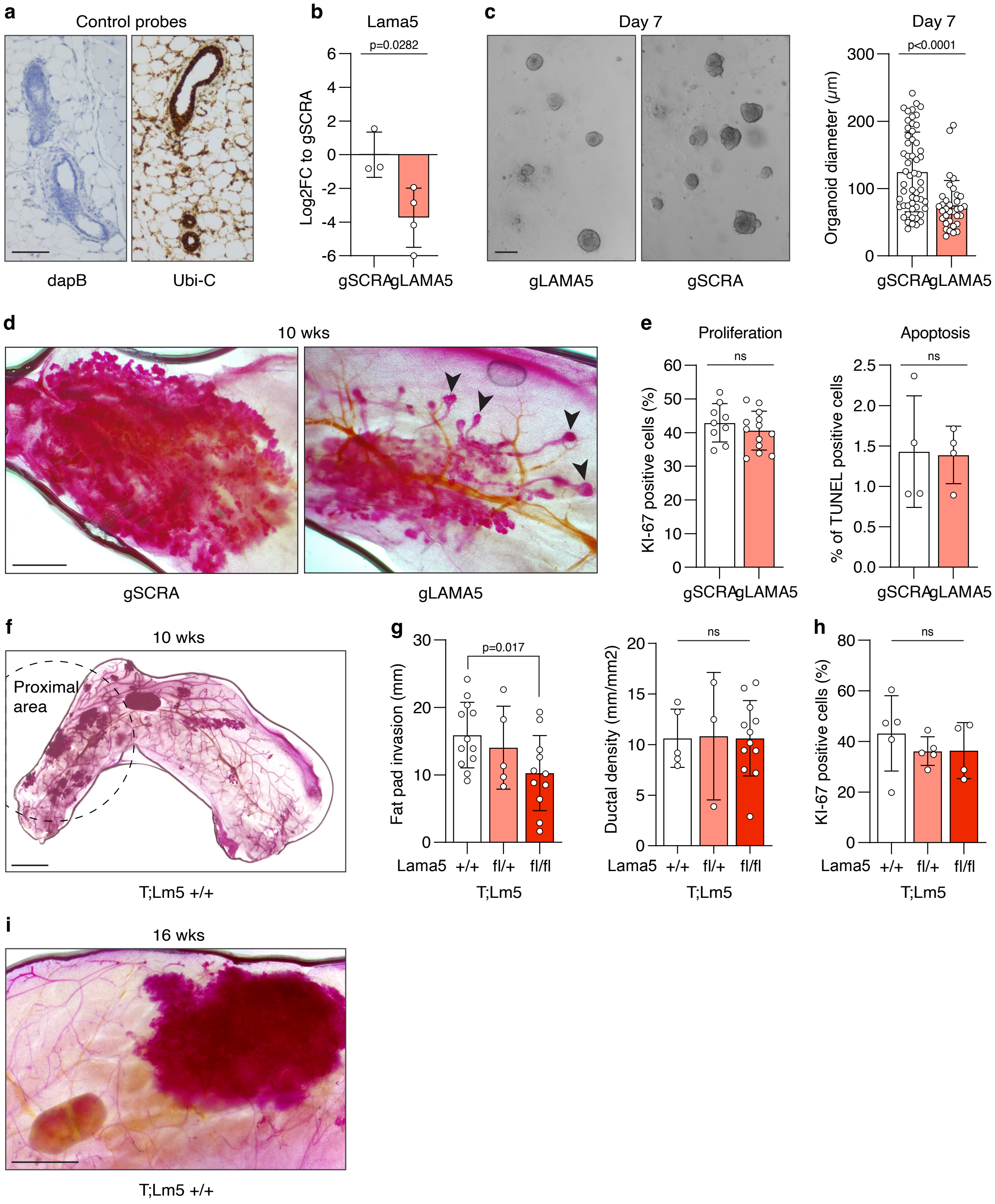


**Supplemental figure 2. Tumour-derived Lama5 is critical for early tumour growth**

**a)** Control ISH stainings of dapB (negative control) and Ubi-C (positive control) in normal mammary gland. Scale bar 20μm. **b)** qPCR analysis comparing LAMA5 expression between MMTV-PyMT hyperplastic cells expressing either LAMA5 or SCRA gDNA. Unpaired t-test was used for statistical analysis. **c)** Representative images of organoids grown from aforementioned cells. Scale bar 200 μm. Graph shows analysis of organoid diameter after 7 days in culture. Statistical analysis was performed with two-tailed Welch's t test. **d)** Representative HE images of mammary glands transplanted with aforementioned cells. Arrow heads denote the normal duct like growth in LAMA5 gDNA glands. Scale bar 1 mm. **e)** Analysis of TUNEL positivity and Ki-67 positivity in the gLAMA5 or gSCRA lesions. Each dot represents one gland analysed. Two-tailed Student's t test was used for statistical testing. **f)** Representative carmine-alumn stained mammary gland with representation of the proximal area used in quantitation. Scale bar is 2 mm. **g)** Quantitation of the length of mammary epithelium growth into the fat pad and ratio of fat pad invasion and ductal length. One-way ANOVA or two-tailed Student's t-test were used for statistical testing. h) Quantitation of Ki-67 positivity in the indicated genotypes. Each dot represents one gland analysed. Statistical testing was performed using One-way ANOVA test. i) Representative HE image of a 16-week-old T;Lm5 mammary gland with a large advanced lesion. Scale bar is 2 mm.
