## Supplemental Figure 3 for "Tumour-derived LAMA5 is critical for tumour initiation and controls progression and phenotype in luminal breast cancer"

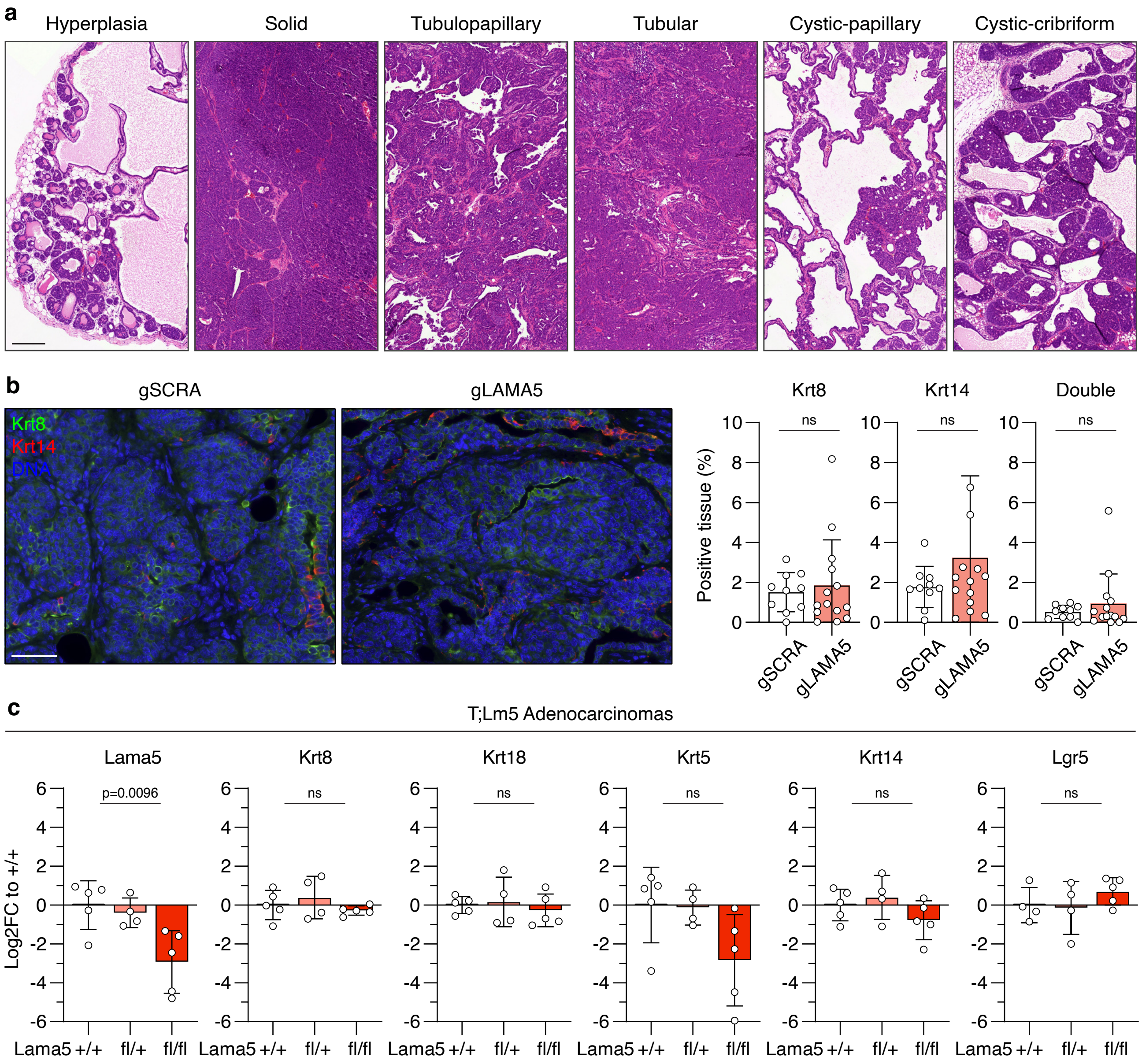


**Supplemental figure 3. Biallelic Lama5 deletion in adenocarcinomas selects for LAMA5-independent growth without altering the tumour subtype**

**a**) Representative HE images of histological subtypes of MMTV-PyMT tumours.Scale bar 200 μm **b)** Representative IF images of transplanted gSCRA and gLAMA5 tumours stained with Krt8 (green) and Krt14 (red) antibodies. Scale bar 50 μm. Graphs show quantiation of Krt8 and Krt14 positivity, as well as double positivity (Krt8 and Krt14) in the lesions. Statistical analysis is performed using two-tailed Welch's t test. Each spot represents 1 tumour analysed. c) qPCR analysis of luminal (Krt8, Krt18) and basal (Krt5, Krt14 and Lgr5) markers analysed in Lama5 low T;Lm5 adenocarcinomas with late Lama5 deletion from indicated genotypes. Each dot represents one carcinoma analysed. Statistical analysis is performed using ordinary one-way ANOVA with Tukey's multiple comparison test.
