## Supplemental Figure 4 for "Tumour-derived LAMA5 is critical for tumour initiation and controls progression and phenotype in luminal breast cancer"

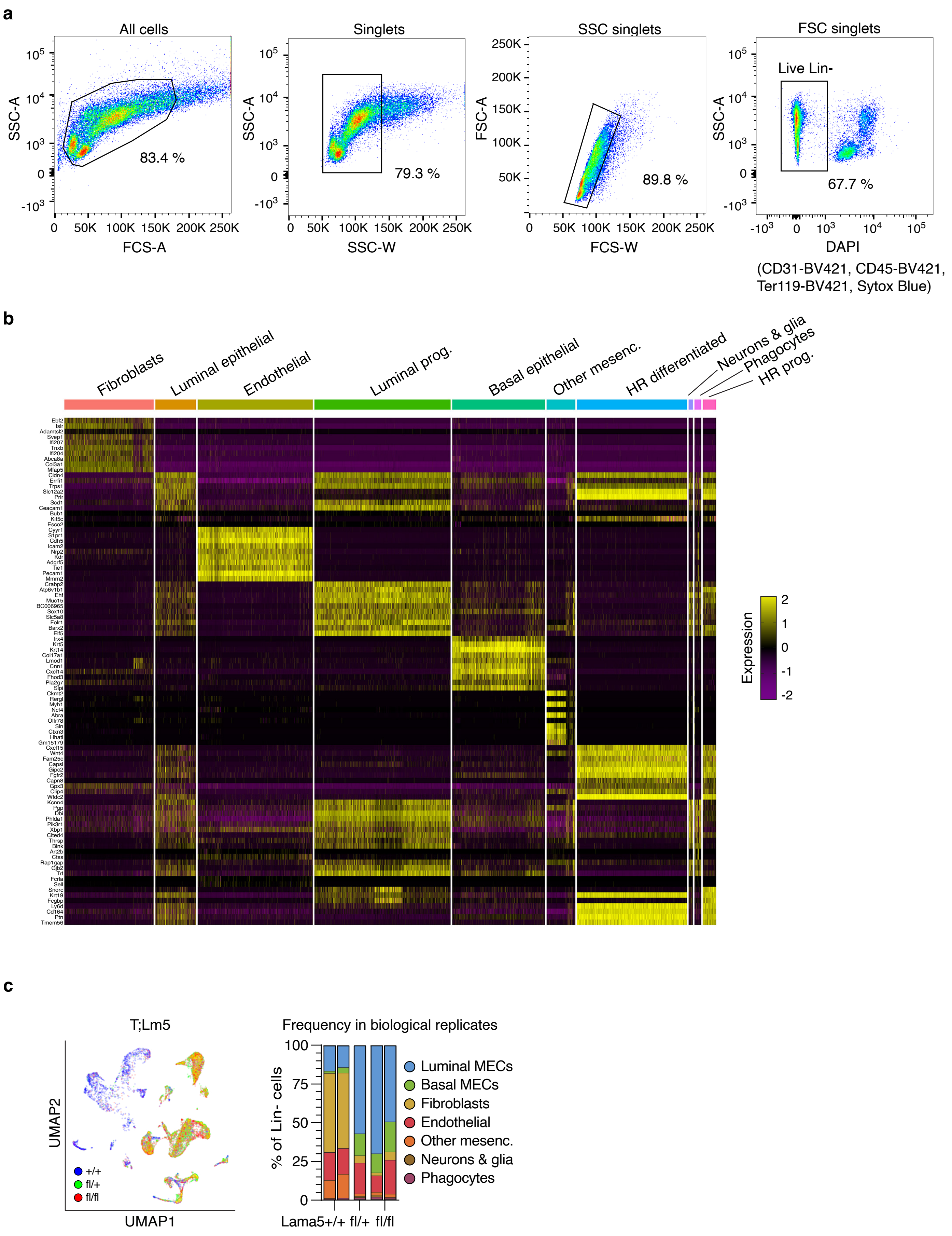


**Supplemental figure 4. Single cell RNA sequencing of Lama5 deficient mammary epithelium**

**a)** Representative FACS plots and gating strategy for collection of live, lineage negative (Lin-) single cells for sc-RNA sequencing. **b)** Heat map showing the top 10 expressed genes contributing to the indicated clusters in the MMTV-PyMT samples. **c)** UMAP showing division of cells by genotype. Graph shows frequency of the clusters identified in each biological T;Lm5 replicate.
