## Supplemental Figure 5 for "Tumour-derived LAMA5 is critical for tumour initiation and controls progression and phenotype in luminal breast cancer"

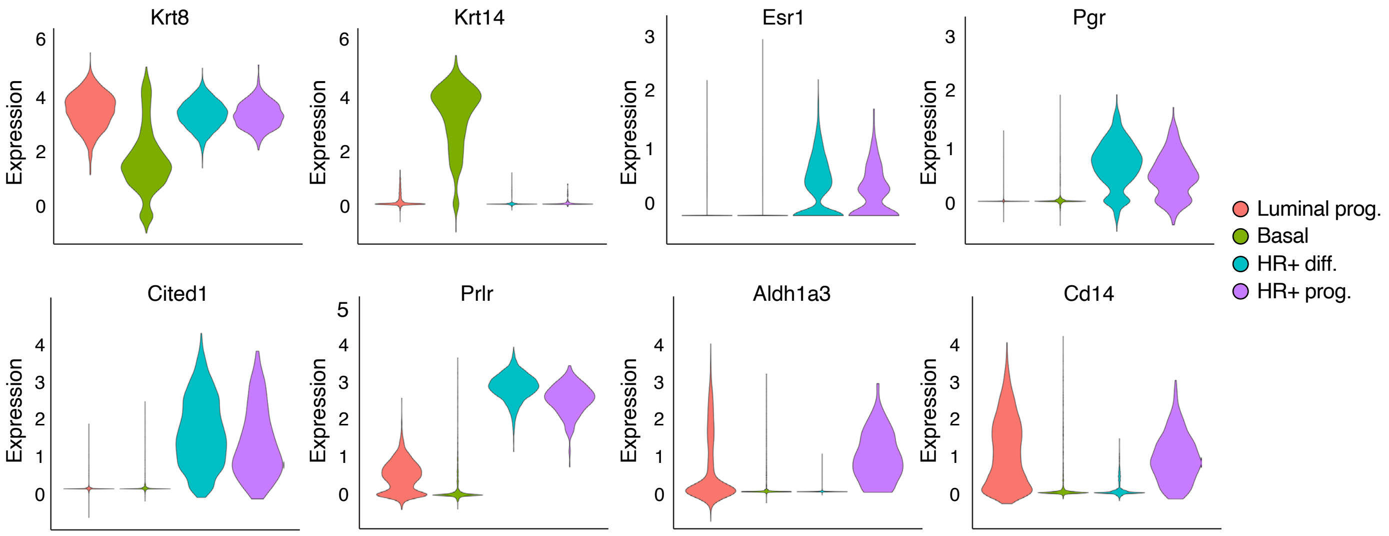


**Supplemental figure 5. LAMA5 deletion from pubertal mammary epithelium and early tumours alters epithelial cell populations**

Violin plots showing the normalized expression of markers previously demonstrated to define the indicated epithelial cell populations (Krt8 - luminal, Krt14 - basal, Esr1, Pgr and Cited1, Prlr - HR+ differentiated and progenitor, Aldh1a3 and Cd14 - progenitor) in main epithelial populations present in MMTV-PyMT samples.
