## Supplemental Figure 6 for "Tumour-derived LAMA5 is critical for tumour initiation and controls progression and phenotype in luminal breast cancer"

**a**

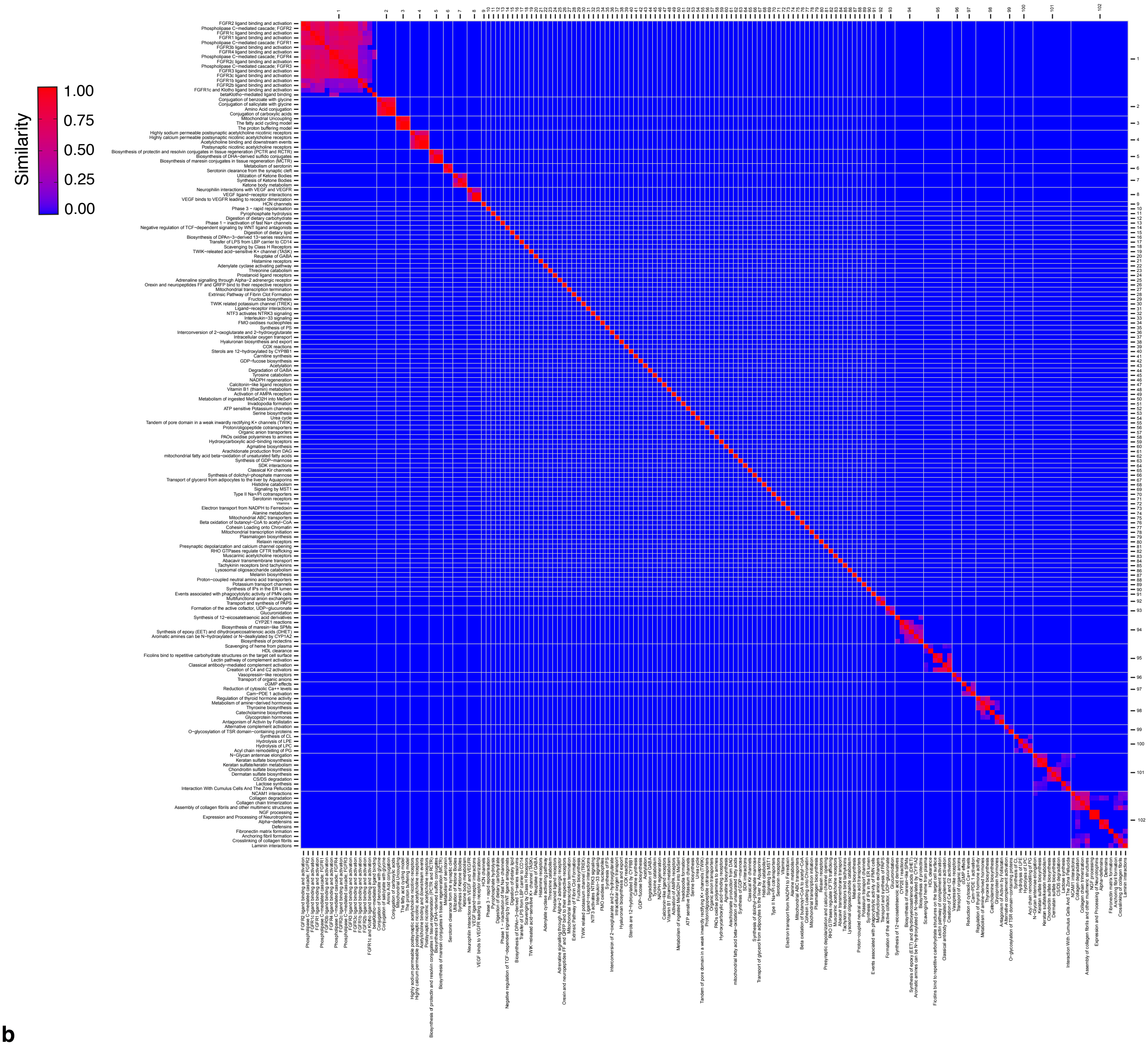


**b**

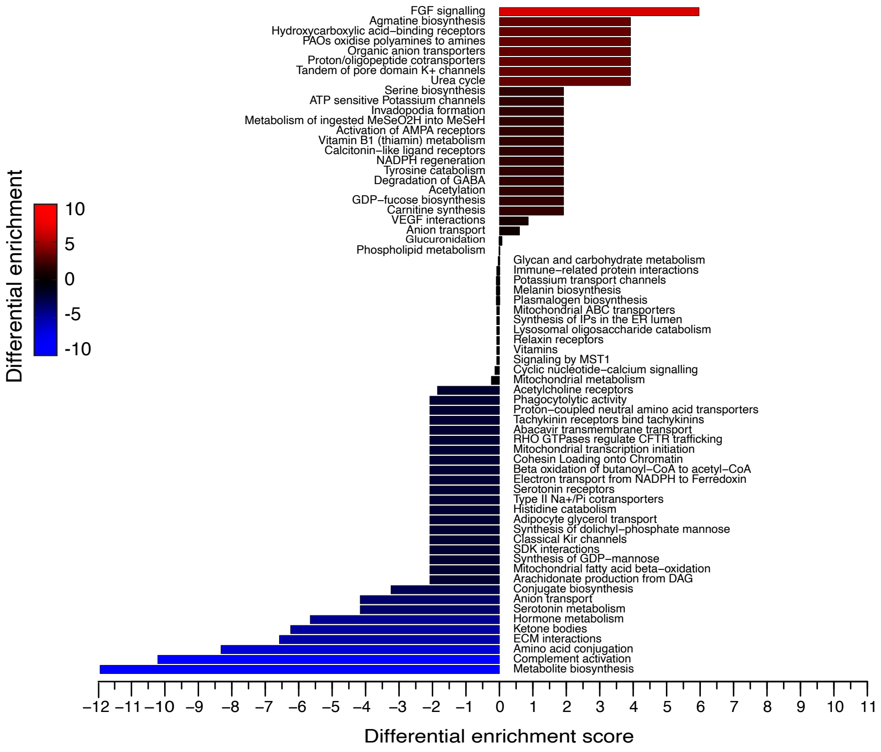


**Supplementary figure 6. Lama5 loss alters gene expression associated with ECM and FGF signalling**

a) Pathway similarity matrix and clustering of pathways enriched in MMTV-PyMT tumor samples, showing similarity between two clusters as gene overlap. Left-bottom axis labels denote the pathway name and top-right axis labels the cluster number. White lines show the pathways arranged to each cluster. **b)** Full list of differential pathway cluster enrichment of Lama5 fl/fl and Lama5 fl/+ tumors compared to Lama5 +/+ control tumors.
