## Supplemental Figure 7 for "Tumour-derived LAMA5 is critical for tumour initiation and controls progression and phenotype in luminal breast cancer"

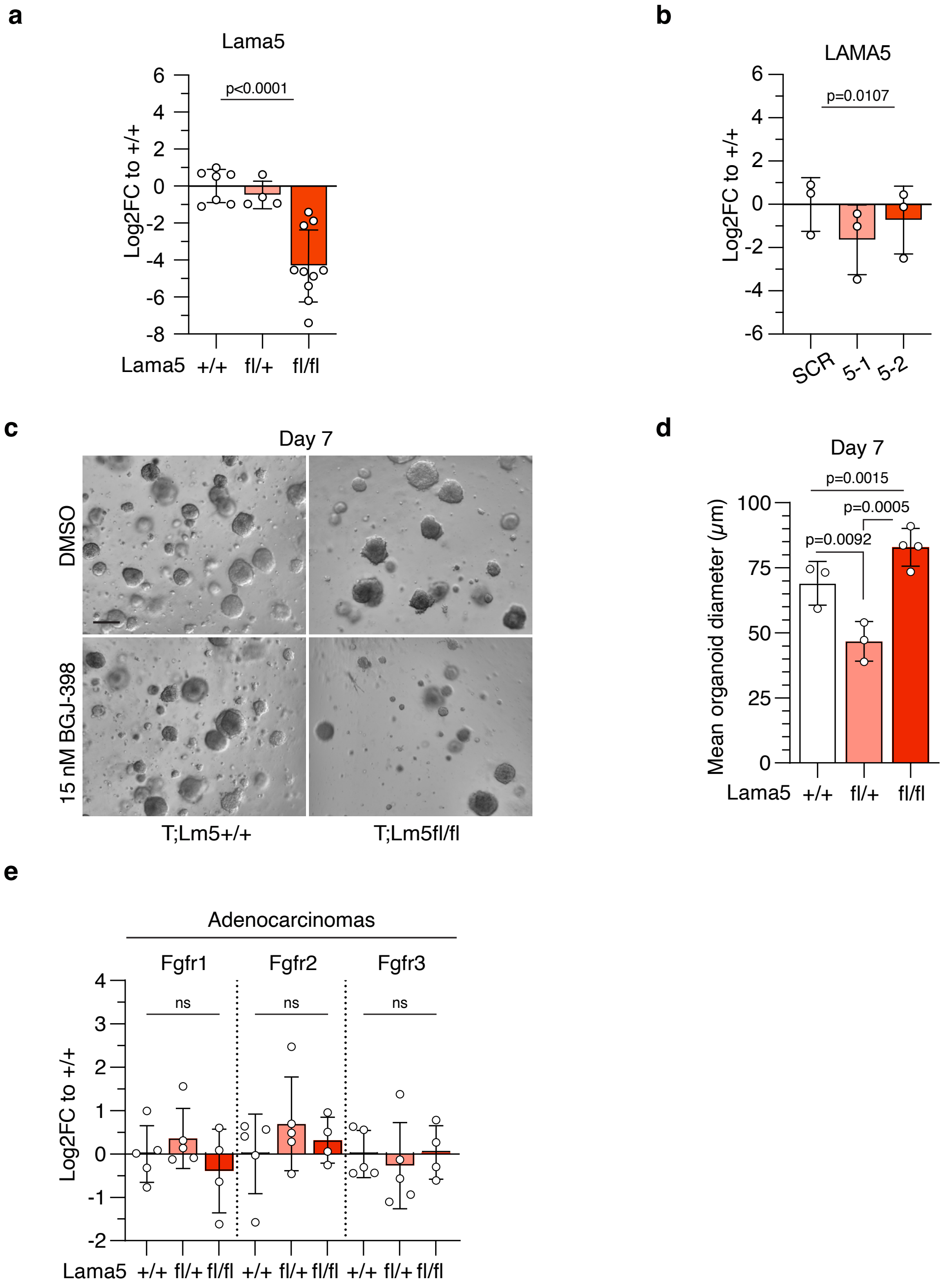


**Supplemental figure 7. Fgfr signaling in Lama5-deficient organoids and tumors**

**a**) qPCR analysis of Lama5 in MECs freshly isolated from T;Lm5fl/+, T;Lm5fl/fl and control T;Lm5+/+ mice. **b**) qPCR analysis of LAMA5 in MCF-7 cells with cells with LAMA5 downregulation (5-1, 5-2) or shCtrl (SCR). Each dot represents a biological replicate. Statistical analysis is performed using ANOVA. **c)** Day 7 organoids isolated from T;Lm5+/+ and T;Lm5fl/fl mice and treated with 15nM FGFRi (BGJ-398) or DMSO starting from day 1 in culture. Scale bar 100 μm. **d**) Quantitation of mean organoid diameter in the DMSO treated organoids at day 7. One dot represents organoids from one mouse. **e)** qPCR analysis of *Fgfr1-3* in T;Lm5 adenocarcinomas of indicated genotypes with Lama5 deletion at 8 weeks of age. Each dot represents 1 tumour. Statistical analysis is performed using ANOVA.
