## Supplemental Figure 1 for "Tumour-derived LAMA5 is critical for tumour initiation and controls progression and phenotype in luminal breast cancer"

| Lama5 genotype | Mouse # | Histological classification* | Tubule formation * | Nuclear pleomorphism | Mitoses per 10 HPF | Sum score | Grade |
|---|---|---|---|---|---|---|---|
| +/+ | 1 | Tubulopapillary carcinoma | 1 | 1 | 1 | 3 | 1 |
| +/+ | 2 | Solid carcinoma | 3 | 3 | 3 | 9 | 3 |
| +/+ | 3 | Solid carcinoma | 2 | 3 | 3 | 8 | 3 |
| +/+ | 4 | Tubular carcinoma | 2 | 2 | 3 | 7 | 3 |
| +/+ | 5 | Tubular carcinoma | 1 | 2 | 1 | 4 | 2 |
| +/+ | 6 | Tubular carcinoma | 1 | 2 | 1 | 4 | 2 |
| +/+ | 7 | Tubular carcinoma | 1 | 1 | 1 | 3 | 1 |
| +/+ | 8 | Tubular carcinoma | 1 | 1 | 1 | 3 | 1 |
| +/+ | 9 | Cystic-cribriform carcinoma | 1 | 2 | 1 | 4 | 2 |
| fl/+ | 10 | Tubular carcinoma | 1 | 1 | 2 | 4 | 2 |
| fl/+ | 11 | Cystic-papillary carcinoma | 1 | 1 | 1 | 3 | 1 |
| fl/+ | 12 | Tubular carcinoma | 3 | 1 | 2 | 6 | 2 |
| fl/+ | 13 | Cystic-papillary carcinoma | 3 | 2 | 3 | 8 | 3 |
| fl/+ | 14 | Solid carcinoma | 2 | 3 | 3 | 8 | 3 |
| fl/+ | 15 | Solid carcinoma | 2 | 3 | 3 | 8 | 3 |
| fl/+ | 15 | Tubulopapillary carcinoma | 2 | 3 | 3 | 8 | 3 |
| fl/+ | 16 | Tubular carcinoma | 1 | 2 | 1 | 4 | 2 |
| fl/+ | 17 | Tubulopapillary carcinoma | 2 | 1 | 2 | 5 | 2 |
| fl/fl | 18 | Tubular carcinoma | 1 | 1 | 2 | 4 | 2 |
| fl/fl | 19 | Tubular carcinoma | 2 | 2 | 3 | 7 | 3 |
| fl/fl | 20 | Solid carcinoma | 3 | 3 | 3 | 9 | 3 |
| fl/fl | 20 | Tubular carcinoma | 1 | 2 | 2 | 5 | 2 |
| fl/fl | 21 | Tubulopapillary carcinoma | 2 | 1 | 1 | 4 | 2 |
| fl/fl | 22 | Tubular carcinoma | 2 | 2 | 1 | 5 | 2 |
| fl/fl | 23 | Solid carcinoma | 3 | 2 | 2 | 7 | 3 |
| fl/fl | 24 | Solid carcinoma | 2 | 3 | 1 | 6 | 2 |

**) according to Goldscmidt et al*

| *Tubule formation* | *Nuclear pleomorphism* | *Mitoses per 10 HPF* | *Grade* |
|---|---|---|---|
| 1 >75 % | 1 Uniform regular nuclei | 1 0-9 | 1 3 |
| 2 10-75% | 2 Moderate degree of variation | 2 10-19 | 2 4-6 |
| 3 <10% | 3 Marked variation | 3 >20 | 3 >7 |

| Lama5 | Grades | n tumors | % of grades |
|---|---|---|---|
| +/+ | 1 | 3 | 33,33333333 |
| +/+ | 2 | 3 | 33,33333333 |
| +/+ | 3 | 3 | 33,33333333 |
| | = | 9 | 100 |

| Lama5 | Grades | n tumors | % of grades |
|---|---|---|---|
| fl/+ | 1 | 1 | 11,11111111 |
| fl/+ | 2 | 4 | 44,44444444 |
| fl/+ | 3 | 4 | 44,44444444 |
| | = | 9 | 100 |

| Lama5 | Grades | n tumors | % of grades |
|---|---|---|---|
| fl/fl | 1 | 0 | 0 |
| fl/fl | 2 | 5 | 62,5 |
| fl/fl | 3 | 3 | 37,5 |
| | = | 8 | 100 |
