## Supplemental Table 2 for "Tumour-derived LAMA5 is critical for tumour initiation and controls progression and phenotype in luminal breast cancer"

| Cluster # | Overlap (Lama5 +/+) | Overlap (Lama5 fl/+) | Overlap (Lama5 fl/fl) | Similarity score | Genes | Gene list | Pathways | Pathway list | Cluster name |
|---|---|---|---|---|---|---|---|---|---|
| **1** | 0,814814815 | 0,111111111 | 0,962962963 | 5,968024682 | 27 | Fgf1 Fgf15 Fgf17 Fgf18 Fgf2 Fgf4 Fgf6 Fgf8 Fgf9 Fgfr4 Fgf23 Fgf20 Fgf16 Klb Fgf5 Galnt3 Fgfr1 Tgfbr3 Gipc1 Fgf10 Fgf22 Kl Fgf7 Fgfbp1 Fgfr2 Fgfbp3 Plcg1 | 16 | betaKlotho-mediated ligand binding, FGFR1 ligand binding and activation, FGFR1b ligand binding and activation, FGFR1c and Klotho ligand binding and activation, FGFR1c ligand binding and activation, FGFR2 ligand binding and activation, FGFR2b ligand binding and activation, FGFR2c ligand binding and activation, FGFR3 ligand binding and activation, FGFR3b ligand binding and activation, FGFR3c ligand binding and activation, FGFR4 ligand binding and activation, Phospholipase C-mediated cascade: FGFR1, Phospholipase C-mediated cascade; FGFR2, Phospholipase C-mediated cascade; FGFR3, Phospholipase C-mediated cascade; FGFR4 | **FGF signalling** |
| **60** | 0 | 1 | 1 | 3,919352858 | 2 | Azin2 Agmat | 1 | Agmatine biosynthesis | Agmatine biosynthesis |
| **59** | 0 | 1 | 1 | 3,919352858 | 2 | Hcar1 Hcar2 | 1 | Hydroxycarboxylic acid-binding receptors | Hydroxycarboxylic acid-binding receptors |
| **57** | 0 | 1 | 1 | 3,919352858 | 9 | Slc17a6 Slc17a1 Slc5a8 Slc17a8 Slc17a5 Slc25a10 Slc25a22 Slc25a18 Slc17a7 | 1 | Organic anion transporters | Organic anion transporters |
| **58** | 0 | 1 | 1 | 3,919352858 | 2 | Paox Smox | 1 | PAOs oxidise polyamines to amines | PAOs oxidise polyamines to amines |
| **56** | 0 | 1 | 1 | 3,919352858 | 3 | Slc15a4 Slc15a1 Slc15a3 | 1 | Proton/oligopeptide cotransporters | Proton/oligopeptide cotransporters |
| **55** | 0 | 1 | 1 | 3,919352858 | 3 | Kcnk1 Kcnk7 Kcnk6 | 1 | Tandem of pore domain in a weak inwardly rectifying K+ channels (TWIK) | **Tandem of pore domain K+ channels** |
| **54** | 0 | 1 | 1 | 3,919352858 | 10 | Asl Arg1 Arg2 Ass1 Slc25a15 Otc Nags Cps1 Nmral1 Slc25a2 | 1 | Urea cycle | Urea cycle |
| **43** | 0 | 0 | 1 | 1,919352858 | 3 | Nat1 Nat2 Nat3 | 1 | Acetylation | Acetylation |
| **49** | 0 | 0 | 1 | 1,919352858 | 3 | Gria1 Gria4 Gria3 | 1 | Activation of AMPA receptors | Activation of AMPA receptors |
| **52** | 0 | 0 | 1 | 1,919352858 | 4 | Kcnj11 Kcnj8 Abcc8 Abcc9 | 1 | ATP sensitive Potassium channels | ATP sensitive Potassium channels |
| **47** | 0 | 0 | 1 | 1,919352858 | 9 | Adm Calca Calcr Iapp Adm2 Ramp1 Ramp2 Calcrl Ramp3 | 1 | Calcitonin-like ligand receptors | Calcitonin-like ligand receptors |
| **41** | 0 | 0 | 1 | 1,919352858 | 3 | Bbox1 Shmt1 Aldh9a1 | 1 | Carnitine synthesis | Carnitine synthesis |
| **44** | 0 | 0 | 1 | 1,919352858 | 2 | Aldh5a1 Abat | 1 | Degradation of GABA | Degradation of GABA |
| **42** | 0 | 0 | 1 | 1,919352858 | 6 | Gmds Gfus Slc35c1 Fcsk Fuom Fpgt | 1 | GDP-fucose biosynthesis | GDP-fucose biosynthesis |
| **51** | 0 | 0 | 1 | 1,919352858 | 4 | Adam12 Adam15 Adam19 Sh3pxd2a | 1 | Invadopodia formation | Invadopodia formation |
| **50** | 0 | 0 | 1 | 1,919352858 | 1 | Txnrd1 | 1 | Metabolism of ingested MeSeO2H into MeSeH | Metabolism of ingested MeSeO2H into MeSeH |
| **46** | 0 | 0 | 1 | 1,919352858 | 1 | Idh1 | 1 | NADPH regeneration | NADPH regeneration |
| **53** | 0 | 1 | 0 | 1,919352858 | 9 | Psph Psat1 Serinc5 Serinc2 Phgdh Serinc3 Srr Serinc1 Serinc4 | 1 | Serine biosynthesis | Serine biosynthesis |
| **45** | 0 | 0 | 1 | 1,919352858 | 5 | Fah Gstz1 Hgd Hpd Tat | 1 | Tyrosine catabolism | Tyrosine catabolism |
| **48** | 0 | 0 | 1 | 1,919352858 | 5 | Thtpa Slc19a2 Tpk1 Slc25a19 Slc19a3 | 1 | Vitamin B1 (thiamin) metabolism | Vitamin B1 (thiamin) metabolism |
| **8** | 0 | 0 | 0,4 | 0,863223429 | 10 | Flt1 Kdr Nrp1 Nrp2 Vegfd Flt4 Pgf Vegfa Vegfb Vegfc | 3 | Neurophilin interactions with VEGF and VEGFR, VEGF binds to VEGFR leading to receptor dimerization, VEGF ligand-receptor interactions | **VEGF interactions** |
| **96** | 0,4 | 0 | 0,666666667 | 0,603581588 | 15 | Avp Avpr2 Oxt Oxtr Avpr1b Avpr1a Slco2b1 Slco4a1 Slco3a1 Slc16a2 Slco4c1 Slco2a1 Slco1a4 Slco1b2 Slco1c1 | 2 | Transport of organic anions, Vasopressin-like receptors | **Anion transport** |
| **93** | 0 | 0 | 0,15 | 0,065805857 | 20 | Ugp2 Ugdh Slc35d1 Ugt2b38 Ugt2b34 Ugt3a1 Ugt2b37 Abhd10 Ugt1a2 Ugt3a2 Ugt2b36 Ugt2b35 Ugt1a7c Ugt1a5 Ugt1a9 Ugt1a1 Ugt2a2 Ugt2b1 Ugt2a1 Ugt1a6a | 2 | Formation of the active cofactor, UDP-glucuronate, Glucuronidation | **Glucuronidation** |
| **100** | 0 | 0,045454545 | 0 | 0,001865809 | 22 | Pla2g4c Crls1 Pla2g4a Pla2g15 Pla2g4b Pla2g4f Pla2g4e Plbd1 Pla2g4d Pla2g1b Pla2r1 Pla2g2a Pla2g2d Pla2g5 Lpcat1 Lpgat1 Pla2g3 Pla2g10 Pla2g2e Pla2g2f Pla2g12a Lpcat4 | 4 | Acyl chain remodelling of PG, Hydrolysis of LPC, Hydrolysis of LPE, Synthesis of CL | **Phospholipid metabolism** |
| **22** | 0 | 0 | 0 | 0 | 9 | Adcy4 Adcy3 Adcy6 Adcy7 Adcy8 Adcy9 Gnal Adcy5 Adcy1 | 1 | Adenylate cyclase activating pathway | Adenylate cyclase activating pathway |
| **25** | 0 | 0 | 0 | 0 | 3 | Adra2a Adra2b Adra2c | 1 | Adrenaline signalling through Alpha-2 adrenergic receptor | Adrenaline signalling through Alpha-2 adrenergic receptor |
| **74** | 1 | 1 | 1 | 0 | 2 | Gpt2 Gpt | 1 | Alanine metabolism | Alanine metabolism |
| **16** | 0 | 0 | 0 | 0 | 1 | Alox5 | 1 | Biosynthesis of DPAn-3-derived 13-series resolvins | Biosynthesis of DPAn-3-derived 13-series resolvins |
| **39** | 0 | 0 | 0 | 0 | 1 | Ptgs1 | 1 | COX reactions | COX reactions |
| **12** | 0 | 0 | 0 | 0 | 10 | Amy2a4 Amy2a3 Amy2a2 Amy2a5 Amy1 Lct Mgam Sis Chit1 Chia1 | 1 | Digestion of dietary carbohydrate | Digestion of dietary carbohydrate |
| **15** | 0 | 0 | 0 | 0 | 6 | Clps Cel Pnliprp1 Pnliprp2 Lipf Pnlip | 1 | Digestion of dietary lipid | Digestion of dietary lipid |
| **28** | 0 | 0 | 0 | 0 | 5 | F10 F3 F7 F9 Tfpi | 1 | Extrinsic Pathway of Fibrin Clot Formation | Extrinsic Pathway of Fibrin Clot Formation |
| **34** | 0 | 0 | 0 | 0 | 3 | Fmo1 Fmo3 Fmo2 | 1 | FMO oxidises nucleophiles | FMO oxidises nucleophiles |
| **29** | 0 | 0 | 0 | 0 | 2 | Akr1b1 Sord | 1 | Fructose biosynthesis | Fructose biosynthesis |
| **9** | 0 | 0 | 0 | 0 | 3 | Hcn1 Hcn2 Hcn3 | 1 | HCN channels | HCN channels |
| **21** | 0 | 0 | 0 | 0 | 4 | Hrh1 Hrh2 Hrh4 Hrh3 | 1 | Histamine receptors | Histamine receptors |
| **38** | 0 | 0 | 0 | 0 | 5 | Has1 Has2 Has3 Abcc5 Cemip | 1 | Hyaluronan biosynthesis and export | Hyaluronan biosynthesis and export |
| **36** | 0 | 0 | 0 | 0 | 3 | L2hgdh Adhfe1 D2hgdh | 1 | Interconversion of 2-oxoglutarate and 2-hydroxyglutarate | **Invadopodia formation** |
| **33** | 0 | 0 | 0 | 0 | 3 | Il1rap Il1rl1 Il33 | 1 | Interleukin-33 signaling | Interleukin-33 signaling |

| | | | | | | | | | |
|---|---|---|---|---|---|---|---|---|---|
| **37** | 0 | 0 | 0 | 0 | 3 | Cygb Mb Ngb | 1 | Intracellular oxygen transport | Intracellular oxygen transport |
| **31** | 0 | 0 | 0 | 0 | 7 | Dhh Gas1 Hhip Ihh Ptch1 Shh Cdon | 1 | Ligand-receptor interactions | Ligand-receptor interactions |
| **27** | 0 | 0 | 0 | 0 | 1 | Mterf1b | 1 | Mitochondrial transcription termination | Mitochondrial transcription termination |
| **83** | 1 | 1 | 1 | 0 | 5 | Chrm1 Chrm3 Chrm4 Chrm5 Chrm2 | 1 | Muscarinic acetylcholine receptors | Muscarinic acetylcholine receptors |
| **14** | 0 | 0 | 0 | 0 | 8 | Dkk1 Lrp5 Lrp6 Dkk4 Dkk2 Kremen2 Sost Kremen1 | 1 | Negative regulation of TCF-dependent signaling by WNT ligand antagonists | |
| **32** | 0 | 0 | 0 | 0 | 2 | Ntf3 Ntrk3 | 1 | NTF3 activates NTRK3 signaling | NTF3 activates NTRK3 signaling |
| **26** | 0 | 0 | 0 | 0 | 8 | Npffr2 Hcrt Qrfp Qrfpr Hcrtr1 Npffr1 Hcrtr2 Npff | 1 | Orexin and neuropeptides FF and QRFP bind to their respective receptors | |
| **13** | 0 | 0 | 0 | 0 | 7 | Kcnd1 Kcnd2 Kcnip3 Kcnd3 Kcnip1 Kcnip4 Kcnip2 | 1 | Phase 1 - inactivation of fast Na+ channels | Phase 1 - inactivation of fast Na+ channels |
| **10** | 0 | 0 | 0 | 0 | 8 | Akap9 Kcna5 Kcnh2 Kcnq1 Kcne2 Kcne3 Kcne4 Kcne5 | 1 | Phase 3 - rapid repolarisation | Phase 3 - rapid repolarisation |
| **81** | 1 | 1 | 1 | 0 | 10 | Cacna1a Cacna1b Cacna2d3 Cacnb1 Cacnb2 Cacnb3 Cacnb4 Cacng2 Cacng4 Cacna2d2 | 1 | Presynaptic depolarization and calcium channel opening | Presynaptic depolarization and calcium channel opening |
| **24** | 0 | 0 | 0 | 0 | 8 | Ptgdr2 Ptgdr Ptger1 Ptger2 Ptger4 Ptgfr Ptgir Tbxa2r | 1 | Prostanoid ligand receptors | Prostanoid ligand receptors |
| **11** | 0 | 0 | 0 | 0 | 3 | Ppa1 Ppa2 Lhpp | 1 | Pyrophosphate hydrolysis | Pyrophosphate hydrolysis |
| **20** | 0 | 0 | 0 | 0 | 3 | Slc6a13 Slc6a1 Slc6a11 | 1 | Reuptake of GABA | Reuptake of GABA |
| **18** | 0 | 0 | 0 | 0 | 3 | Stab2 Sparc Apob | 1 | Scavenging by Class H Receptors | Scavenging by Class H Receptors |
| **40** | 0 | 0 | 0 | 0 | 2 | Cyp8b1 Ptgis | 1 | Sterols are 12-hydroxylated by CYP8B1 | Sterols are 12-hydroxylated by CYP8B1 |
| **35** | 0 | 0 | 0 | 0 | 2 | Ptdss1 Ptdss2 | 1 | Synthesis of PS | Synthesis of PS |
| **23** | 0 | 0 | 0 | 0 | 3 | Rida Sds Sdsl | 1 | Threonine catabolism | Threonine catabolism |
| **17** | 0 | 0 | 0 | 0 | 2 | Cd14 Lbp | 1 | Transfer of LPS from LBP carrier to CD14 | Transfer of LPS from LBP carrier to CD14 |
| **30** | 0 | 0 | 0 | 0 | 3 | Kcnk2 Kcnk4 Kcnk10 | 1 | TWIK related potassium channel (TREK) | TWIK related potassium channel (TREK) |
| **19** | 0 | 0 | 0 | 0 | 2 | Kcnk3 Kcnk9 | 1 | TWIK-releated acid-sensitive K+ channel (TASK) | TWIK-releated acid-sensitive K+ channel (TASK) |
| **101** | 0,037037037 | 0 | 0,037037037 | -0,02389545 | 81 | B4galt1 Lalba Slc2a1 Adam2 Ovgp1 Zp1 Zp2 Zp3 Adam25 Adam21 Adam30 Hyal5 Bcan Cspg4 Bgn Ncan Dcn Dse Cspg5 Dsel Ust Chst14 Chpf2 Csgalnact1 Chsy1 Chst3 Chst11 Chst12 Chst7 Chst9 Chst13 Chpf Chst15 Csgalnact2 Chsy3 Mgat4b Mgat5 St6gal1 St3gal4 St8sia2 St8sia3 St8sia6 Mgat4a B4galt2 B4galt5 B4galt4 B4galt6 B4galt3 Mgat4c Hyal3 Arsb Hexa Hexb Hyal1 Ids B4gat1 Acan Prelp Fmod Kera Lum Ogn St3gal3 St3gal1 St3gal2 B3gnt7 B3gnt4 Omd B3gnt2 Chst2 St3gal6 Chst5 Slc35d2 B3gnt3 Chst1 Glb1 Glb1l2 Galns Glb1l3 Glb1l Gns | 8 | Chondroitin sulfate biosynthesis, CS/DS degradation, Dermatan sulfate biosynthesis, Interaction With Cumulus Cells And The Zona Pellucida, Keratan sulfate biosynthesis, Keratan sulfate/keratin metabolism, Lactose synthesis, N-Glycan antennae elongation | **Glycan and carbohydrate metabolism** |
| **99** | 0,111111111 | 0 | 0 | -0,067304303 | 36 | Cfd C3 Gzmm Cfp Spon2 Adamts7 Adamts6 Adamts1 Sema5b Thsd4 Adamts18 Thsd7b Adamts2 Thbs2 Adamts20 Adamts10 Sbspon Adamtsl4 Adamts17 Spon1 Adamts15 Adamts5 Adamts12 Adamts19 Adamts4 Adamtsl3 Adamts16 Adamts13 Adamts3 Thsd7a B3glct Thsd1 Adamtsl5 Adamtsl1 Adamtsl2 Pofut2 | 2 | Alternative complement activation, O-glycosylation of TSR domain-containing proteins | **Immune-related protein interactions** |
| **87** | 1 | 1 | 0 | -0,080647142 | 5 | Dct Oca2 Tyr Tyrp1 Slc45a2 | 1 | Melanin biosynthesis | Melanin biosynthesis |
| **75** | 1 | 1 | 0 | -0,080647142 | 3 | Abcb7 Abcb6 Abcb8 | 1 | Mitochondrial ABC transporters | Mitochondrial ABC transporters |
| **79** | 1 | 1 | 0 | -0,080647142 | 3 | Gnpat Dhrs7b Agps | 1 | Plasmalogen biosynthesis | Plasmalogen biosynthesis |
| **89** | 1 | 1 | 0 | -0,080647142 | 3 | Kcnj10 Kcnj16 Kcnj1 | 1 | Potassium transport channels | Potassium transport channels |
| **86** | 1 | 0 | 1 | -0,080647142 | 4 | Manba Man2b1 Man2b2 Man2c1 | 1 | Lysosomal oligosaccharide catabolism | Lysosomal oligosaccharide catabolism |
| **80** | 1 | 0 | 1 | -0,080647142 | 6 | Rxfp2 Rln3 Insl5 Rxfp3 Rxfp4 Rxfp1 | 1 | Relaxin receptors | Relaxin receptors |
| **69** | 1 | 0 | 1 | -0,080647142 | 5 | Mst1 Hpn Mst1r Spint1 Spint2 | 1 | Signaling by MST1 | Signaling by MST1 |
| **90** | 1 | 0 | 1 | -0,080647142 | 1 | Minpp1 | 1 | Synthesis of IPs in the ER lumen | Synthesis of IPs in the ER lumen |
| **72** | 1 | 0 | 1 | -0,080647142 | 6 | Cyp24a1 Cyp26a1 Cyp27b1 Cyp26b1 Cyp2r1 Cyp26c1 | 1 | Vitamins | Vitamins |
| **97** | 0,56 | 0 | 0,56 | -0,135487199 | 25 | Sri Slc8a2 Slc8a3 Atp2a1 Atp2a2 Atp2b2 Calm1 Calm2 Calm3 Slc8a1 Atp2b3 Atp2b4 Atp2a3 Atp2b1 Pde1a Pde1b Pde1c Itpr1 Irag1 Pde9a Prkg1 Pde2a Pde10a Pde11a Pde5a | 3 | Cam-PDE 1 activation, cGMP effects, Reduction of cytosolic Ca++ levels | **Cyclic nucleotide-calcium signalling** |
| **3** | 1 | 0 | 1 | -0,241941427 | 6 | Slc25a14 Pm20d1 Ucp1 Ucp2 Ucp3 Slc25a27 | 3 | Mitochondrial Uncoupling, The fatty acid cycling model, The proton buffering model | **Mitochondrial metabolism** |
| **4** | 0,461538462 | 0 | 0 | -1,853029044 | 13 | Chrnb4 Chrna3 Chrna4 Chrnb2 Chrnd Chrne Chrnb3 Chrna5 Chrna2 Chrna1 Chrna6 Chrna7 Chrna9 | 4 | Acetylcholine binding and downstream events, Highly calcium permeable postsynaptic nicotinic acetylcholine receptors, Highly sodium permeable postsynaptic acetylcholine nicotinic receptors, Postsynaptic nicotinic acetylcholine receptors | **Acetylcholine receptors** |
| **84** | 1 | 0 | 0 | -2,080647142 | 3 | Slc22a1 Slc22a2 Slc22a3 | 1 | Abacavir transmembrane transport | Abacavir transmembrane transport |
| **61** | 1 | 0 | 0 | -2,080647142 | 5 | Daglb Mgll Dagla Abhd6 Abhd12 | 1 | Arachidonate production from DAG | Arachidonate production from DAG |
| **76** | 1 | 0 | 0 | -2,080647142 | 3 | Acads Hadh Echs1 | 1 | Beta oxidation of butanoyl-CoA to acetyl-CoA | Beta oxidation of butanoyl-CoA to acetyl-CoA |
| **65** | 1 | 0 | 0 | -2,080647142 | 3 | Kcnj12 Kcnj2 Kcnj14 | 1 | Classical Kir channels | Classical Kir channels |
| **77** | 1 | 0 | 0 | -2,080647142 | 9 | Pds5b Smc3 Rad21 Stag1 Stag2 Wapl Smc1a Nipbl Mau2 | 1 | Cohesin Loading onto Chromatin | Cohesin Loading onto Chromatin |
| **73** | 1 | 0 | 0 | -2,080647142 | 3 | Fdx1 Fdxr Fdx2 | 1 | Electron transport from NADPH to Ferredoxin | Electron transport from NADPH to Ferredoxin |
| **91** | 1 | 0 | 0 | -2,080647142 | 2 | Mpo Lpo | 1 | Events associated with phagocytolytic activity of PMN cells | **Phagocytolytic activity** |
| **68** | 1 | 0 | 0 | -2,080647142 | 6 | Ftcd Hal Hdc Uroc1 Cammt1 Amdhd1 | 1 | Histidine catabolism | Histidine catabolism |

| | | | | | | | | | |
|---|---|---|---|---|---|---|---|---|---|
| **62** | 1 | 0 | 0 | -2,080647142 | 3 | Acadm Eci1 Decr1 | 1 | mitochondrial fatty acid beta-oxidation of unsaturated fatty acids | **Mitochondrial fatty acid beta-oxidation** |
| **78** | 1 | 0 | 0 | -2,080647142 | 3 | Tfb2m Polrmt Tfam | 1 | Mitochondrial transcription initiation | Mitochondrial transcription initiation |
| **88** | 1 | 0 | 0 | -2,080647142 | 2 | Slc36a1 Slc36a2 | 1 | Proton-coupled neutral amino acid transporters | Proton-coupled neutral amino acid transporters |
| **82** | 1 | 0 | 0 | -2,080647142 | 3 | Rhoq Cftr Gopc | 1 | RHO GTPases regulate CFTR trafficking | RHO GTPases regulate CFTR trafficking |
| **64** | 1 | 0 | 0 | -2,080647142 | 2 | Sdk2 Sdk1 | 1 | SDK interactions | SDK interactions |
| **71** | 1 | 0 | 0 | -2,080647142 | 11 | Htr1a Htr1b Htr1d Htr1f Htr2a Htr2b Htr2c Htr4 Htr5a Htr6 Htr7 | 1 | Serotonin receptors | Serotonin receptors |
| **66** | 1 | 0 | 0 | -2,080647142 | 3 | Dpm1 Dpm2 Dpm3 | 1 | Synthesis of dolichyl-phosphate mannose | Synthesis of dolichyl-phosphate mannose |
| **63** | 1 | 0 | 0 | -2,080647142 | 3 | Mpi Pmm1 Pmm2 | 1 | Synthesis of GDP-mannose | Synthesis of GDP-mannose |
| **85** | 1 | 0 | 0 | -2,080647142 | 5 | Tac1 Tac2 Tacr1 Tacr2 Tacr3 | 1 | Tachykinin receptors bind tachykinins | Tachykinin receptors bind tachykinins |
| **67** | 1 | 0 | 0 | -2,080647142 | 2 | Aqp7 Aqp9 | 1 | Transport of glycerol from adipocytes to the liver by Aquaporins | **Adipocyte glycerol transport** |
| **70** | 1 | 0 | 0 | -2,080647142 | 2 | Slc34a3 Slc34a2 | 1 | Type II Na+/Pi cotransporters | Type II Na+/Pi cotransporters |
| **5** | 1 | 0 | 0,5 | -3,241941427 | 2 | Ltc4s Gstm4 | 3 | Biosynthesis of DHA-derived sulfido conjugates, Biosynthesis of maresin conjugates in tissue regeneration (MCTR), Biosynthesis of protectin and resolvin conjugates in tissue regeneration (PCTR and RCTR) | **Conjugate biosynthesis** |
| **6** | 1 | 0 | 0 | -4,161294285 | 3 | Aldh2 Maoa Slc6a4 | 2 | Metabolism of serotonin, Serotonin clearance from the synaptic cleft | **Serotonin metabolism** |
| **92** | 1 | 0 | 0 | -4,161294285 | 12 | Slc26a3 Slc26a2 Slc26a6 Slc26a7 Slc26a1 Slc26a4 Slc5a12 Slc26a11 Slc26a9 Slc35b3 Papss1 Papss2 | 2 | Multifunctional anion exchangers, Transport and synthesis of PAPS | **Anion transport** |
| **98** | 0,666666667 | 0 | 0 | -5,655921903 | 24 | Dbh Ddc Pnmt Th Dio3 Asmt Aanat Cga Dio1 Dio2 Duox2 Tph2 Tph1 Tpo Iyd Duox1 Fshb Inha Inhba Inhbb Inhbc Inhbe Fst Fstl3 | 6 | Antagonism of Activin by Follistatin, Catecholamine biosynthesis, Glycoprotein hormones, Metabolism of amine-derived hormones, Regulation of thyroid hormone activity, Thyroxine biosynthesis | **Hormone metabolism** |
| **7** | 1 | 0 | 0 | -6,241941427 | 11 | Acat1 Hmgcl Hmgcs2 Hmgcll1 Acss3 Bdh2 Bdh1 Aacs Oxct2b Oxct2a Oxct1 | 3 | Ketone body metabolism, Synthesis of Ketone Bodies, Utilization of Ketone Bodies | **Ketone bodies** |
| **102** | 0,503703704 | 0 | 0 | -6,576685279 | 135 | Defa30 Prss3l Defa35 Defa41 Defa40 Defa27 Defa32 Defa37 Defa34 AY761185 Defa36 Try5 Prss1 Art1 Cd4 Defa31 Defa25 Defa3 Prss2 Prss3 Try4 Defa17 Defa39 Defa22 Try10 Prss1l Defa23 Defa24 Defa28 Defa26 Defa38 Defa21 Defa42 Defa43 Defa20 Ccr6 Defb1 Tlr1 Tlr2 Defb14 Defb19 Defb36 Defb21 Defb48 Defb28 Defb4 Defb42 Defb43 Defb25 Defb18 Defb47 Defb30 Bmp1 Col4a1 Col4a2 Col4a5 Col1a1 Col1a2 Lox Loxl1 Loxl3 Pcolce Tll2 Loxl4 Pxdn Col4a6 Loxl2 Col7a1 Col16a1 Col10a1 Col11a1 Col11a2 Col12a1 Col13a1 Col14a1 Col15a1 Col17a1 Col18a1 Col19a1 Col2a1 Col3a1 Col5a1 Col5a2 Col6a1 Col6a2 Col8a1 Col9a1 Col9a3 Col26a1 Col28a1 Col23a1 Col6a6 Col8a2 Col27a1 Col5a3 Col6a5 Col22a1 Col24a1 Col20a1 Col25a1 Ctsb Ctsl Ctss Mmp13 Mmp3 Mmp9 Mmp20 Ncam1 St8sia2 St8sia4 Ngf Furin Pcsk5 Pcsk6 Itga6 Itgb1 Lama4 Nid1 Nid2 Megf11 Fn1 Itga5 Ceacam1 Ctsd Ctsk Mmp12 Mmp10 Mmp11 Mmp14 Mmp15 Mmp2 Mmp8 Tmprss6 Phykpl Mmp1a | 12 | Alpha-defensins, Anchoring fibril formation, Assembly of collagen fibrils and other multimeric structures, Collagen chain trimerization, Collagen degradation, Crosslinking of collagen fibrils, Defensins, Expression and Processing of Neurotrophins, Fibronectin matrix formation, Laminin interactions, NCAM1 interactions, NGF processing | **ECM interactions** |
| **2** | 1 | 0 | 0 | -8,32258857 | 6 | Glyat Acsm1 Acsm2 Acsm4 Acsm5 Glyatl3 | 4 | Amino Acid conjugation, Conjugation of benzoate with glycine, Conjugation of carboxylic acids, Conjugation of salicylate with glycine | **Amino acid conjugation** |
| **95** | 1 | 0 | 0,189189189 | -10,21361258 | 37 | C1qa C1qb C1qc Crp Igll1 C1s2 C1ra Fcna Fcnb Masp1 Masp2 Mbl2 Colec10 Colec11 Hbb-bs Hbb-bt Apol7c Alb Ambp Apoa1 Hp Hpx Jchain Lrp1 Apol9a Apol8 Apol10a Apol7b Apol10b Apol11b Apol11a Apol7e Apol9b Apol7a Cd163 Hdlbp Scarb1 | 6 | Classical antibody-mediated complement activation, Creation of C4 and C2 activators, Ficolins bind to repetitive carbohydrate structures on the target cell surface, HDL clearance, Lectin pathway of complement activation, Scavenging of heme from plasma | **Complement activation** |
| **94** | 1 | 0 | 0,045454545 | -11,93842831 | 22 | Cyp1a2 Cyp2a12 Cyp2a4 Cyp2e1 Cyp2f2 Cyp2d22 Cyp2c66 Cyp2c65 Cyp2s1 Alox15 Cyp1a1 Lta4h Alox12 Alox12b Gpx1 Gpx2 Aloxe3 Gpx4 Cyp3a11 Cyp1b1 Cyp2j6 Ephx2 | 6 | Aromatic amines can be N-hydroxylated or N-dealkylated by CYP1A2, Biosynthesis of maresin-like SPMs, Biosynthesis of protectins, CYP2E1 reactions, Synthesis of 12-eicosatetraenoic acid derivatives, Synthesis of epoxy (EET) and dihydroxyeicosatrienoic acids (DHET) | **Metabolite biosynthesis** |
